# Development of FAP-targeted theranostics discovered by next-generation sequencing-augmented mining of a novel immunized VNAR library

**DOI:** 10.1101/2025.01.13.632555

**Authors:** Gihan S. Gunaratne, Joseph P. Gallant, Kendahl L. Ott, Payson L. Broome, Sasha Celada, Jayden L. West, Jason C. Mixdorf, Eduardo Aluicio-Sarduy, Jonathan W. Engle, Eszter Boros, Labros Meimetis, Joshua M. Lang, Shuang G. Zhao, Reinier Hernandez, David Kosoff, Aaron M. LeBeau

## Abstract

Cancer-associated fibroblasts (CAFs) in the stroma of solid tumors promote an immunosuppressive tumor microenvironment (TME) that drives resistance to therapies. The expression of the protease fibroblast activation protein (FAP) on the surface of CAFs has made FAP a target for development of therapies to dampen immunosuppression. Relatively few biologics have been developed for FAP and none have been developed that exploit the unique engagement properties of Variable New Antigen Receptors (VNARs) from shark antibodies. As the smallest binding domain in nature, VNARs cleverage unique geometries and recognize epitopes conventional antibodies cannot. By directly immunizing a nurse shark with FAP, we created a large anti-FAP VNAR phage display library. This library allowed us to identify a suite of anti-FAP VNARs through traditional biopanning and also by an *in silico* approach that did not require any prior affinity-based enrichment *in vitro*. We investigated four VNAR-Fc fusion proteins for theranostic properties and found that all four recognized FAP with high affinity and were rapidly internalized by FAP-positive cells. As a result, the VNAR-Fc constructs were effective antibody-drug conjugates *in vitro* and were able to localize to FAP-positive xenografts *in vivo*. Our findings establish VNAR-Fc constructs as a versatile platform for theranostic development that could yield innovative cancer therapies targeting the TME.

## Introduction

The immunosuppressive microenvironment of solid tumors hinders the effectiveness of immune checkpoint inhibitors and novel cellular therapies at treating cancer^1,2^. Several cell types contribute to an immunosuppressive tumor microenvironment (TME) that promotes tumor growth, metastasis, and drug resistance^3,4^. One of the most abundant cell types in the TME are cancer-associated fibroblasts (CAFs)^5^. Existing as heterogeneous populations within the TME, pro-tumorigenic CAFs play a salient role in immunosuppression by directly recruiting immunosuppressive myeloid cells and inhibiting the function of cytotoxic lymphocytes^6–8^. CAFs expressing the cell surface serine protease fibroblast activation protein (FAP) are highly pro-tumorigenic and are directly associated with the recruitment of immunosuppressive myeloid cells^9–11^. Previously, using a microfluidics model, we demonstrated that the direct elimination of FAP-positive CAFs by an anti-FAP antibody-drug conjugated (ADC) resulted in upregulation of proinflammatory genes, secretion of proinflammatory cytokines, and alterations in the immune microenvironment^12^. Our observations and others showing that eliminating FAP-positive CAFs in immunocompetent mouse models increased CD8+ tumor infiltrating lymphocytes suggest that FAP-targeted therapies represent a strategy for enhancing the immune anti-tumor response^12,13^.

The expression of FAP has been reported in the TME of nearly every type of solid tumor^14,15^. Under normal physiologic conditions, FAP expression is restricted to tissues undergoing wound healing and embryogenesis with little to no expression in healthy adult tissues^16^. The disease specificity of FAP has spurred interest in the development of diagnostics and therapeutics for cancer. Once heralded as the next billion-dollar theranostic target, countless small-molecules and peptides have been developed targeting the active site of FAP^17–20^. These include quinoline-based FAP inhibitors (FAPIs), the cyclic peptide FAP-2286, and the peptidyl boronic acid inhibitor PNT6555. Both FAPIs and FAP-2286 have been used to image more than two dozen different cancer types by PET^21,22^. Several clinical trials are currently underway investigating these molecules for radioligand therapy using the β-emitting radionuclides ^90^Y and ^177^Lu. Several antibody-based therapies targeting FAP have been developed including the humanized antibody sibrotuzumab which showed no therapeutic efficacy in a phase II trial.^23^ Since sibrotuzumab, several bispecific antibodies and immunocytokines have been investigated in the clinic^24–26^. These biologics were either abandoned or met with limited success^24,27–29^. Several CAR-T cell therapies for FAP have been explored in preclinical models; however, none have been translated into the clinic^30^. Thus far, no FAP-targeted therapies have been approved by the FDA and no therapies have made it past Phase II.

Conventional antibodies consisting of heavy and light chain binding domains have been used for decades in the clinic as targeted therapies. There are disadvantages associated with the use of conventional antibodies including poor pharmacokinetics, difficulty engineering, and high production costs. The complementarity determining regions (CDRs) of conventional antibodies also prefer planar epitopes limiting their modes of engagement. The structural and functional limitations of conventional antibodies have increased interest in alternative targeting scaffolds including single-domain antibodies. Antigen binding domains from camelid antibodies, also known as variable-heavy-heavy (VHH) domains or nanobodies, have been utilized for applications ranging from nuclear medicine to opioid use disorder^31,32^. Less explored as targeting vectors are Variable New Antigen Receptors (VNARs) which are the single-chain binding domains of shark antibodies. At a molecular weight of 11kDa, a VNAR is smaller than a human single-chain variable fragment (scFv, 25kDa), and a camelid VHH domain (15kDa). Though VNARs only have two CDRs, the protruding geometry of the CDRs coupled with the presence of two hypervariable loops in the framework regions allows VNARs to bind cryptic epitopes inaccessible to conventional human and camelid antibodies.

Through the direct immunization of a juvenile male nurse shark with human FAP (hFAP), we created a high diversity FAP-biased VNAR phage display library. Using a traditional biopanning strategy to identify functional clones, we discovered three unique VNARs (H4, H15, and H17) that recognized hFAP from the library. Next-generation sequencing (NGS) found a richly diverse library biased towards the Type II VNAR subtype. By performing sequence alignment analyses of our NGS data in parallel with NGS data from an immunized library of an unrelated immunogen as a control dataset, we were able to identify an entirely unique functional clone (NGS2405) through an *in silico* approach without any prior affinity-based enrichment *in vitro*. As bivalent human IgG1 Fc fusion proteins, our VNAR-Fc constructs were able to bind recombinant FAP and FAP-positive cells though biolayer interferometry (BLI) and flow cytometry. Using two separate complementary methods, we determined that the VNAR-Fc constructs were rapidly and specifically internalized by FAP-positive cells. Due to internalization, the VNAR-Fc constructs were potent ADCs when coupled to an anti-mitotic payload. Three of the VNAR-Fc constructs demonstrated high localization to FAP-positive xenografts by PET imaging suggestive of potential theranostic applications *in vivo*. Our study documents an important proof-of-concept showing for the first time that potent VNAR targeting vectors for a cancer-associated antigen can be identified via the direct immunization of a nurse shark. The VNAR-Fc constructs reported here represent a suite of first-in-class agents that could have a profound impact on how the TME is targeted in solid tumors.

## Results

### Shark immunization yields high-affinity anti-FAP VNARs

To identify anti-FAP VNARs, a juvenile male nurse shark was immunized with recombinant hFAP using complete Freund’s adjuvant. Subsequent booster immunizations were performed every two weeks, either subcutaneously using incomplete Freund’s adjuvant or intravenously via the caudal vein with free hFAP to ensure the antigen is presented in a near native conformation **(Figure 1A-C)**. Blood samples were collected prior to beginning the immunization program (pre-bleed), immediately before each antigen administration, and two weeks after the final immunization. Mobilization of an anti-FAP immune response was monitored by screening shark plasma for the presence of anti-FAP convalescent IgNARs using biolayer interferometry (BLI). Purified recombinant hFAP was biotinylated and immobilized on Octet streptavidin (SAX) biosensors and exposed to diluted plasma collected from each blood draw. No binding was detected in naïve plasma isolated from the pre-bleed sample, whereas samples collected throughout the immunization displayed a time-dependent increase in hFAP binding with a pronounced spike observed in samples collected after the third immunization **(Figure 1D-E)**. Plasma collected from the final time point failed to produce a response in control biosensors lacking hFAP (no bait). The immune repertoire was captured by cloning the VNAR sequences present in buffy coat samples isolated from blood draws. To mitigate biased representation of high-frequency clones that may result from T-cell clonal expansion at later time points, RNA was isolated from samples collected on weeks 6 and 8. VNAR-encoding sequences were amplified and used to generate a FAP-biased VNAR phage-display library (∼8 x 10^8^ cfu), which was screened against hFAP. After a single round of biopanning, 83 out of 192 selected clones were found to be strong FAP binders by ELISA **(Figure 1F)**. Sanger sequencing of ‘hit’ clones revealed 17 unique VNARs, with one cluster of 14 sequences sharing a high degree (>95%) of amino acid sequence conservation, a second grouping of 2 highly homologous VNARs, and a final discrete VNAR sequence (H17) **(Figure 1G, Supplemental Figure 1).**

**Figure 1.**
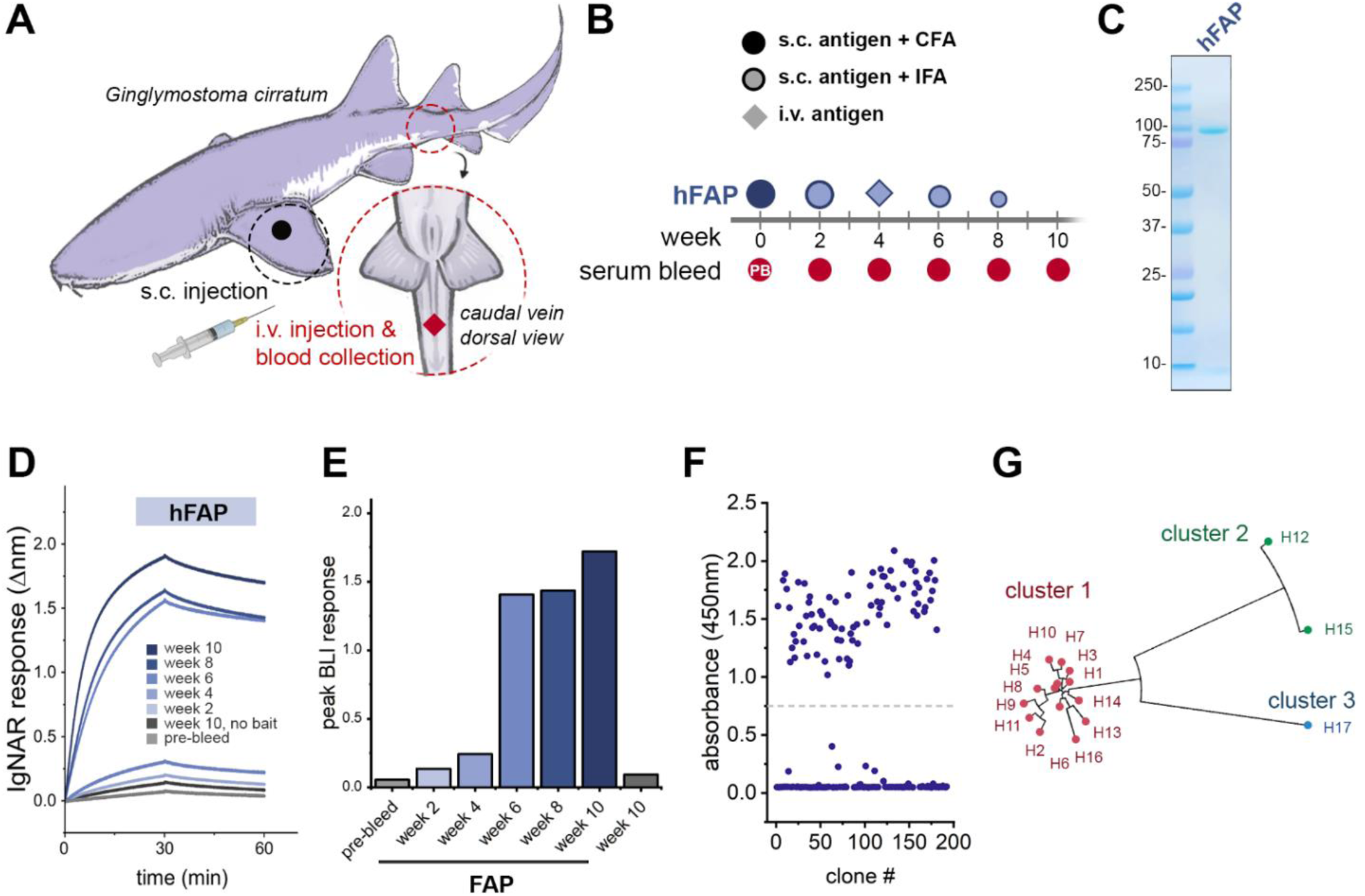
Immunization of a live nurse shark and identification of anti-FAP VNARs. **A)** Schematic of sites used for subcutaneous (s.c.) or intravenous (i.v.) delivery of immunogens and blood collection. **B)** Illustration of the time course, injection sites, adjuvants used, and blood sample collection schedule throughout the FAP immunization program. **C)** SDS-PAGE and Coomassie staining of purified recombinant human FAP (hFAP) protein used for immunization. **D)** Biolayer interferometry (BLI) sensorgram from a representative experiment demonstrating the mobilization of an anti-FAP immune response after hFAP immunization. Diluted plasma samples (1:200) collected from the indicated time points were screened against biosensors loaded with immobilized hFAP for the presence of convalescent anti-hFAP IgNARs. Control sensors were exposed to plasma from week 10 in the absence of hFAP ligand. **E)** Quantification of convalescent anti-FAP IgNAR response in the indicated time points, data represents peak Δnm after 30min of dissociation. **F)** 196 clones were screened by ELISA for the production of anti-hFAP VNARs after a single round of biopanning by phage display. A threshold absorbance (OD450nm) of 0.75 was used for the identification of positive clones. **G)** Unrooted phylogenetic tree illustrating the relative sequence homology of positive anti-FAP VNAR clones, sequences sharing >90% sequence homology are depicted with the same color.

### Next-generation sequencing of the FAP-immunized VNAR phage display library

To understand the diversity and composition of the VNAR clones arising from the nurse shark immune response, we conducted NGS analysis on the library. VNAR-encoding sequences were digested out of phagemid samples and analyzed by forward and reverse paired-end sequencing. Of 5.3×10^6^ reads, 1.2×10^6^ reads encoded full-length VNARs. cDNA sequences were translated to amino acids, and repeated sequences were collated and ranked based on prevalence within the dataset **(Figure 2A)**. A small portion of unique VNAR sequences were highly abundant with nearly 1×10^4^ repeats detected, however, the vast majority of sequences detected were fewer than 10 times (99.5%) or only detected with a single read (90.9%), supportive of a richly diverse population of VNARs in the library. To characterize the makeup of the FAP-immunized library, we analyzed the distribution of VNAR subtypes, prevalence of cysteine residues, and length of CDR3 loops of all unique VNAR sequences **(Figure 2B-D)**. The presence of non-canonical cysteine residues in positions 21C, 28C, 34C and 82C, which constrain VNAR tertiary structure through disulfide bonding, are key determinants of VNAR subtype, and by extension, the geometric flexibility of the paratopic loops **(Supplemental Fig. 2)**. Type I VNARs are characterized by a CDR3 that is held in close apposition with HV2 due to disulfide bonding between non-canonical cysteines in framework regions (FR) 2 and 4 with paired cysteines in CDR3. Type II VNARs exploit the lack of CDR3 cysteine pairs to form a probing CDR3 domain that can access clefts and deep motifs. Type III VNARs are structurally like Type II but have restricted CDR3 diversity and sequence conservation in CDR1. Lastly, Type IV VNARs are the most structurally flexible as they only have canonical cysteine residues. The subtype distribution of the FAP-immunized library was heavily biased towards Type II VNARs **(Figure 2B)**. The total number of cysteine residues present in each sequence displayed a modest biphasic pattern driven by cysteine richness within the CDR3 domains **(Figure 2C)**. VNAR CDR3 lengths were found to have a 1.5x interquartile range of 5-25 amino acids **(Figure 2D)** with outliers having hyper-elongated CDR3 loops of over 80 amino acids in length, reminiscent of reports of ultralong ‘stalk and knob’ antibodies from bovine species.^33^

**Figure 2.**
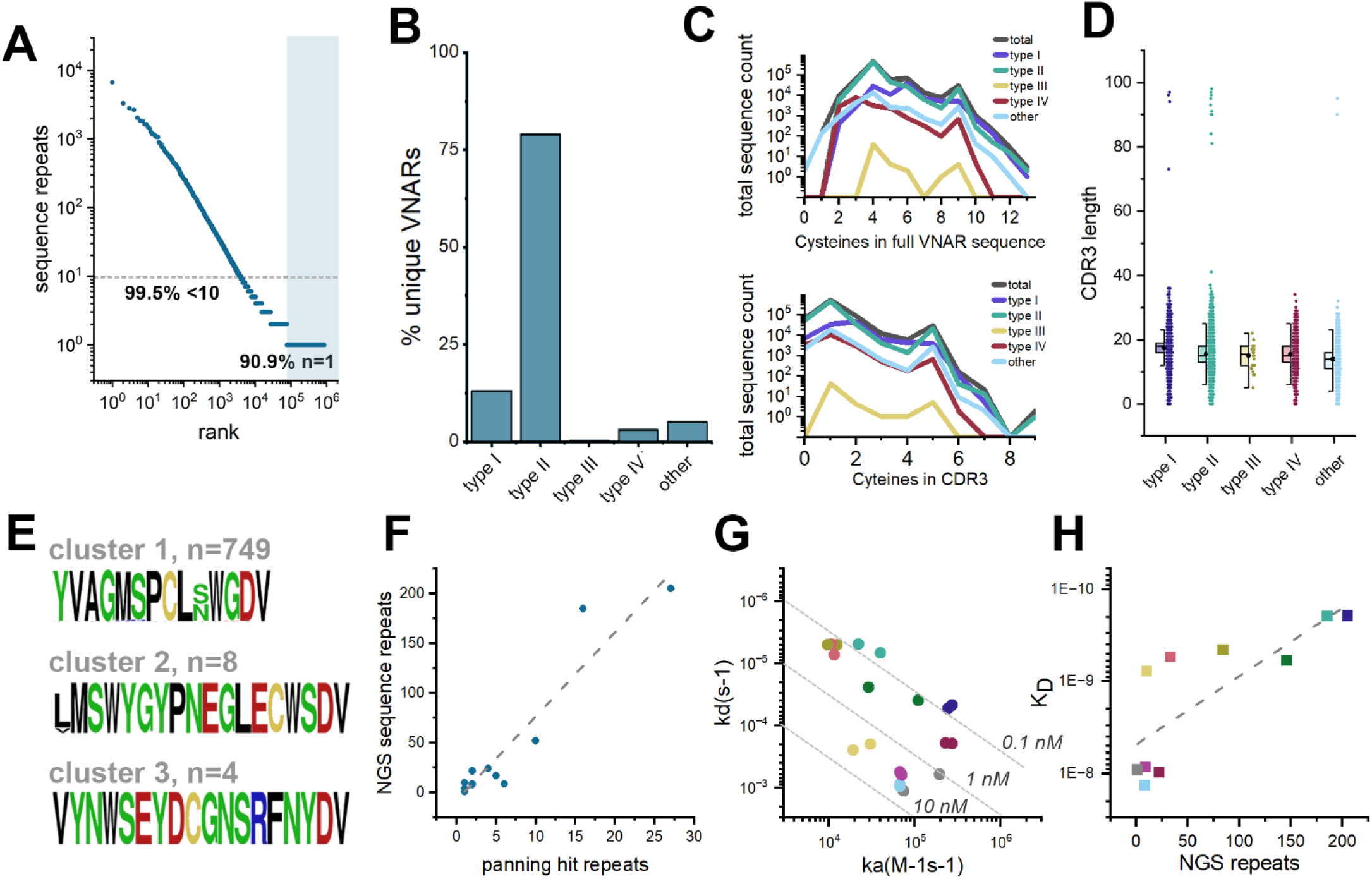
Next generation sequencing of hFAP-immunized phagemid library and validation of anti-FAP VNARs. **A)** hFAP-immunized VNAR phagemid library was analyzed by MiSeq, sequences were ranked based on the prevalence of repeats. Scatter plot represents the rank-ordered distribution of sequence repeats, shown on a log-scale. Dashed line and shaded region indicate clones that are present in the sequencing dataset less than 10 times (99.5%) or a single time (90.9%), respectively. **B)** VNAR subtype distribution among all unique sequences. **C)** Prevalence of cysteine residues in unique full length VNAR sequences (top) or CDR3s of unique VNARs (bottom), per VNAR subtype. **D)** Number of amino acids in the CDR3 of VNARs by subtype. Data is presented as a column scatter, overlaid with a box plot reporting 25-75% percentile (box), mean (closed circle), and median (line). **E)** Sequence logos of the CDR3s of unique VNAR clones present in NGS dataset and share >90% sequence homology with hit anti-FAP VNARs identified by phage display. Polar amino acids (GSTYQN) are green; basic amino acids (KRH) are blue; acidic amino acids (DE) are red; hydrophobic amino acids (AVLIPWFM) are black; cysteines (C) are yellow. **F)** Pearson correlation of the number of sequence repeats per clone, detected by Sanger sequencing of hit clones after phagemid biopanning versus the number of identical sequences detected by NGS of the phagemid library. **G)** Isoaffinity plot of all 10 anti-FAP VNAR clones from biopanning clade 1 which were also detected by NGS. Data points are color coded by clone ID. **H)** Pearson correlation of the number of sequence repeats detected by NGS versus the measured affinity of 10 anti-FAP VNAR clones.

We next investigated how the anti-FAP VNARs identified by phage display compared to VNARs defined by NGS. CDR3 sequences of each of the three VNAR clusters identified by biopanning were used to identify all VNARs in the NGS dataset with a similar clonotype (>85% CDR3 sequence homology). This yielded 749 unique VNARs similar to biopanning cluster 1, 8 unique VNAR sequences belonging to cluster 2, and only 4 unique VNARs homologous to cluster 3 (H17) **(Figure 2E)**. These results were consistent with the relative proportion of VNARs present in each phylogenetic cluster identified by biopanning (**Figure 1G**). These findings also suggest an intuitive relationship between the prevalence of VNAR clones present in the immunized library and the likelihood of identifying the same clone by biopanning. This is supported by a Pearson correlation analysis (R^2^=0.951) of the number of sequence repeats per clone, as detected by Sanger sequencing of hit clones after biopanning, plotted against the number of identical sequences detected by NGS of the phagemid library **(Figure 2F)**. Since the prevalence of antibody sequences within an immune repertoire is partly a function of the ability of the antibody to engage the antigen, we next predicted that anti-FAP VNARs that were highly abundant in the NGS dataset would display a higher affinity for hFAP than low abundance VNAR sequences. Ten VNARs sequences that were abundant in the NGS dataset and also identified by biopanning were grafted onto a human IgG1 Fc domain and expressed as VNAR-Fc fusion proteins. After purification, their affinities for hFAP were determined by BLI **(Figure 2G)**. The clones screened showed subtle variation in their sequences, largely accounted for in the HV4 and FW5 domains (**Supplemental Fig. 1**), however, log-scale changes in affinity were measured with clone H4 emerging as the highest affinity VNAR from biopanning cluster 1. Plotting the raw number of sequence repeats in the NGS dataset against the measured affinity for hFAP for each clone revealed a positive trendline. An R^2^ value of 0.792 indicates that these two variables are not linearly interdependent, however, among the clones identified by biopanning, those that were most highly abundant in the NGS dataset also had the highest affinity for hFAP **(Figure 2H)**.

We next decided to see if an *in silico* approach could be employed to identify anti-FAP VNARs by screening the most abundant VNAR sequences from the NGS dataset. VNAR-Fc constructs of the top seven most prevalent VNAR sequences with unique CDR3s were expressed and screened for their ability to engage hFAP using the newly validated anti-FAP H4-Fc as a positive control **(Figure 3A)**. Despite their high prevalence in the VNAR library, all of the antibodies failed to bind hFAP, suggesting that sequence prevalence cannot be used as the sole metric for the identification of functional antibodies. To more accurately identify antibodies with functional specificity, we performed sequence alignment analyses using sequencing data from the FAP VNAR library in parallel with NGS data from an immunized library of an unrelated immunogen (∼18% sequence homology) as a control dataset. The control dataset contained a nearly identical number of full-length VNAR reads as the FAP-immunized library dataset, with less than 0.2% of sequences being shared among both NGS datasets **(Figure 3B).** To simplify sequence alignment analyses, we constrained alignments to only CDR3 sequences of identical length, and from the top 2000 most prevalent VNARs from both libraries. Since we already experimentally identified a clade of functional anti-FAP VNARs with a CDR3 length of 14 amino acids (**Figure 1G, 2E-H)**, we focused initial efforts on VNARs with CDR3s of 14 amino acids to instill confidence in this approach. Despite the near-absent degree of sequence commonality between the two libraries (**Figure 3B**), sequence alignment analysis of this mixed pool of CDR3s from both libraries yielded only 4 clades with greater than 30 antibodies that were unique to the FAP library (FAP 14aa clades 1-4) **(Figure 3C).** Encouragingly, one of these clades (FAP 14aa clade 3) consisted of the clonotype that defines the dominant cluster of anti-FAP VNARs that were experimentally identified by phage display (**Figure 1G**). Within each clade, the full-length VNAR sequence of the clone with the highest NGS prevalence was expressed as VNAR-Fc fusion proteins, and screened for binding to hFAP. **(Figure 3D)**. As anticipated, clone H5-Fc (NGS131), identified by phage display, showed a robust anti-FAP binding response. An additional 2 of the 3 novel clonotypes displayed FAP binding, albeit with less favorable kinetics **(Figure 3D).** Nonetheless, this improved the hit-rate when compared to testing clones based on library prevalence alone (**Figure 3A**) and provided the impetus for further screening. We next performed a similar analysis with the top 2000 most prevalent antibodies with a CDR3 length of 12 amino acids from both library datasets **(Figure 3E)**, which enabled us to resolve an additional 4 clades (FAP 12aa clades 1-4) that were specifically present in the FAP-immunized library. The most prevalent antibody sequence within each identified clade was then expressed as a VNAR-Fc fusion protein and tested for FAP binding. This resulted in the identification of clones NGS812 and NGS31 as novel anti-FAP clonotypes. Due to its favorable association kinetics, clone NGS812 was prioritized for further investigation. Phylogenetic analysis of the 77 unique VNAR sequences in the NGS dataset that shared >85% CDR3 sequence homology with NGS812 yielded 5 discrete nodes **(Figure 3G)**. The most prevalent clones from each discrete node, NGS2405, NGS10865, NGS812, NGS2582, and NGS2132, were expressed and tested for FAP binding. All 5 clones displayed varying degrees of FAP binding, with NGS2405 emerging as VNAR clone with the highest affinity for FAP **(Figure 3H)**.

**Figure 3.**
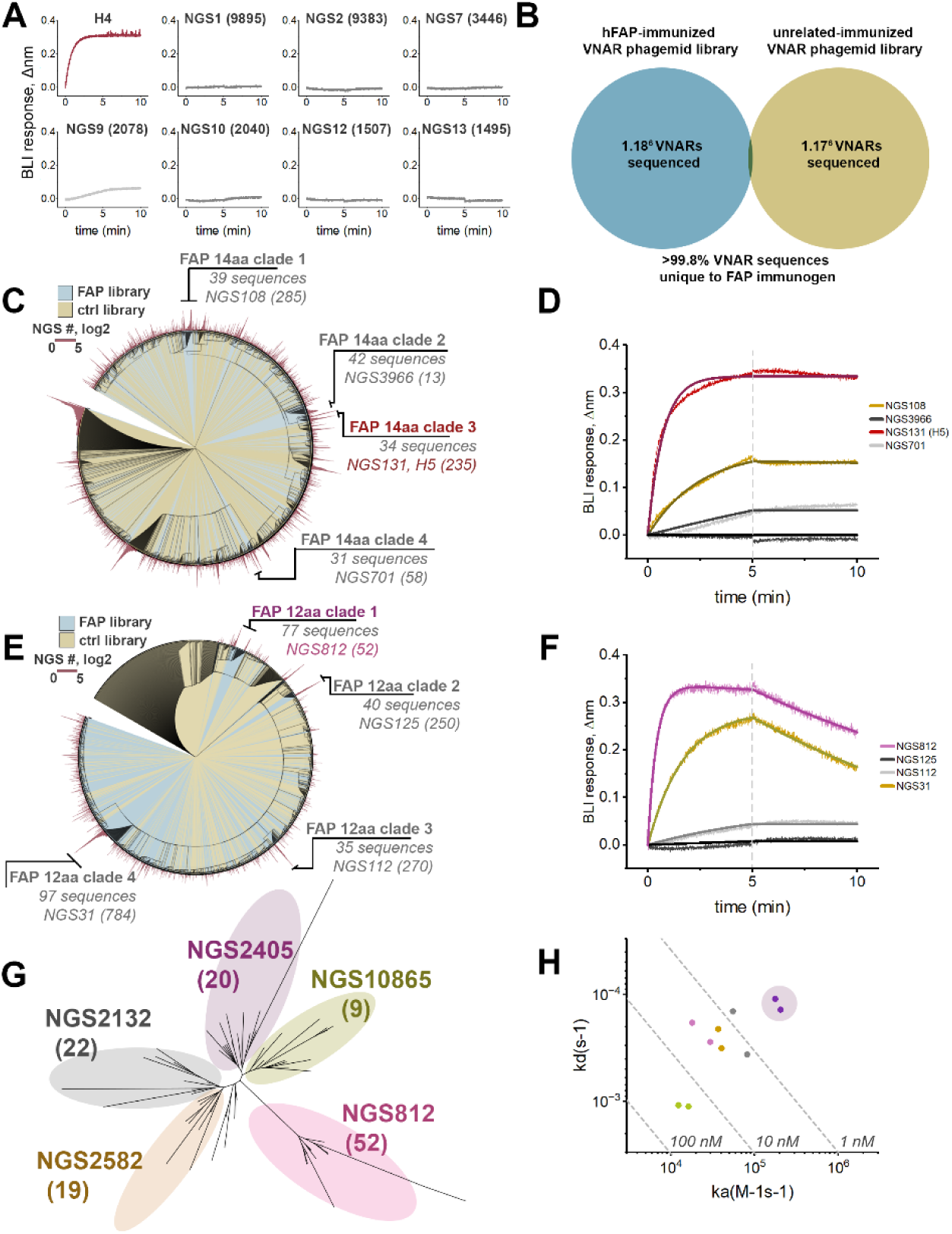
Identification of novel anti-FAP VNAR clones using NGS datasets. **A)** The top 7 most prevalent VNAR sequences with unique CDR3s were screened for anti-hFAP binding by BLI, anti-FAP VNAR H4 was used as a positive control. Clone IDs represent the sequence ‘rank’ as described in Figure 2A, with the number of sequence repeats shown in parentheses. **B)** Venn diagram of sequence overlap between NGS dataset derived from sequencing of FAP-immunized VNAR phagemid library compared to a VNAR phagemid library immunized against a discrete unrelated immunogen. **C)** Circular phylogenetic tree of the top 2000 most prevalent 14-residue long CDR3s from the FAP-immunized NGS dataset (blue) and control dataset (yellow), overlaid with a circularized bar graph (red) representing the number of sequence repeats for each node. Clades with ≥30 unique sequences are shown, along with the most prevalent clone ID and the number of sequencing repeats. **D)** BLI sensorgram of experiment screening the most prevalent VNAR (500nM) from each clade in (C) against biosensors with immobilized hFAP. **E)** Circular phylogenetic tree of the top 2000 most prevalent 12-residue long CDR3s from the FAP-immunized NGS dataset (blue) and control dataset (yellow), overlaid with a circularized bar graph (red) representing the number of sequence repeats for each node. Clades with ≥30 unique sequences are shown, along with the most prevalent clone ID and the number of sequencing repeats. **F)** BLI sensorgram of experiment screening the most prevalent VNAR (500nM) from each clade in (E) against biosensors with immobilized hFAP. **G)** Unrooted phylogenetic tree of all unique sequences in the FAP-immunized phagemid library NGS dataset with a CDR3 length of 12aa and >90% CDR3 homology compared to clone NGS812. Distinct clusters are color coded with most prevalent clone within each cluster shown, along with the number of repeats in the NGS dataset. **H)** Iso-affinity plot of putative anti-FAP VNARs identified in (G). The most prevalent VNAR sequence from each cluster in (G) was screened against hFAP by BLI. The highest affinity clone, NGS2405, is depicted with purple shading.

### *In vitro* characterization of anti-FAP VNAR Fcs

Using relative affinity for hFAP as a means of prioritization, VNARs H4, H15, H17 and NGS2405 were selected as lead representatives of their respective clonotypes. Each lead VNAR was cloned into an expression vector encoding the Fc region of human IgG_1_, expressed, and purified **(Supplemental Fig. 3)**. Dissociation constants (K_D_) of each VNAR-Fc for hFAP were determined by BLI, and were measured to range from 14 pM to 1.3 nM **(Figure 4A)**. Since cross-reactivity with murine FAP is an essential characteristic for validating anti-FAP therapeutic agents in endogenous FAP-expressing mouse models, we determined the affinity of each VNAR-Fc for mouse FAP (mFAP). Both H4-Fc and NGS2405-Fc bound mFAP, while H15-Fc and H17-Fc failed to recognize mFAP **(Figure 4B)**. As FAP is a member of the prolyl protease family, the constructs were then tested against human dipeptidyl peptidase IV (hDPP-IV), a serine protease that serves as the closest related protein to hFAP. All four lead VNAR-Fcs failed to bind hDPP-IV, demonstrating target specificity for FAP **(Supplemental Fig. 4).** K_D_ values of each construct for each of the antigens described are collated in **Table 1**. Next, cross-competition antibody binding assays were conducted to assess epitope overlap. It was revealed that H4-Fc and NGS2405-Fc each recognized discrete epitopes on FAP, while H15-Fc and H17-Fc competed for a third epitope, consistent with their shared inability to bind mFAP **(Figure 4C)**. None of the VNAR-Fc constructs were found to inhibit the proteolytic activity of FAP using either small dipeptide or nonameric fluorogenic peptide substrates **(Figure 4D-G)**. This suggests that their epitopes are distal to the active site and do not occlude substrate binding or non-covalently inhibit proteolysis.

**Figure 4.**
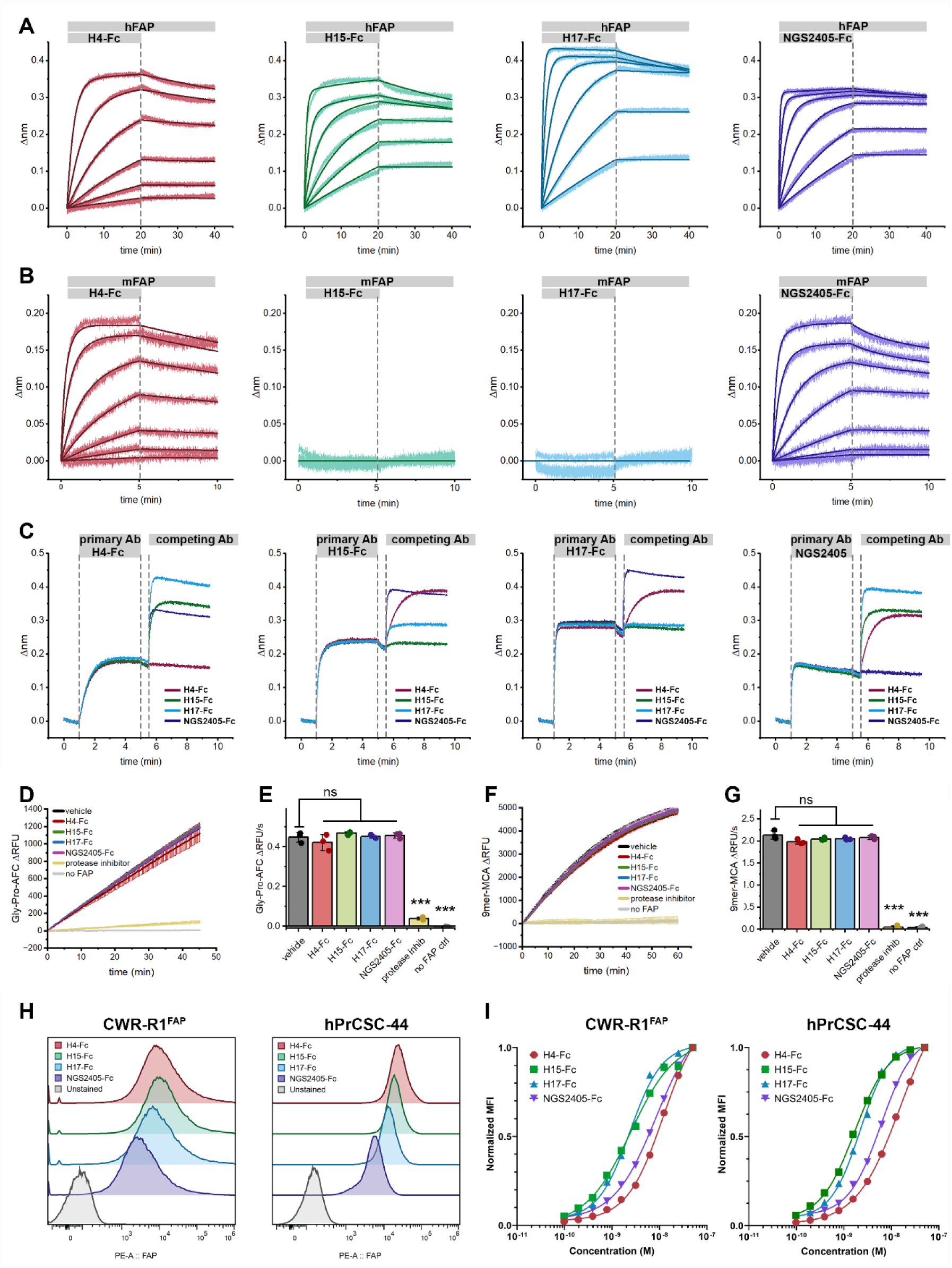
*In vitro* characterization of lead anti-FAP VNAR-Fc constructs. **A-B)** BLI sensorgrams of sensors loaded with hFAP (A) or mFAP (B), and monitored during exposure to serially diluted antibody analytes (300nM-0.412nM), followed by dissociation in assay buffer. Data represents raw BLI responses (thin lines) and fitted curves (bold lines) from a representative experiment. **C)** BLI sensorgrams from antibody cross-competition epitope binning experiments, wherein biosensors loaded with hFAP are exposed to a saturating concentration (1µM) of the indicated primary antibody, followed by exposure to a competing secondary antibody (1µM). **D-G)** kinetic traces (D, F) and cumulative quantification (E, G) of proteolytic activity of recombinant hFAP (0.3nM) in the presence of the indicated VNAR-Fc (1µM) using 1µM of either Ac-Gly-Pro-AFC (D-E) or MCA-Glu-Arg-Gly-Glu-Thr-Gly-Pro-Ser-Gly-Dnp (‘9mer’, F-G) fluorogenic substrates. **H)** Validation of membrane bound FAP expression in R1-CWR_FAP_ and hPrCSC-44 cell lines by flow cytometry. Cells were stained using a fixed concentration of VNAR-Fc (50nM) and detected using a anti-IgG1-phycoerythrin secondary (5µg/mL) . Samples were compared to an unstained cell control. **I)** Dose response curves of R1-CWR_FAP_ and hPrCSC-44 cell lines using several staining concentrations of VNAR-Fc antibodies assessed by flow cytometry. p-values, ***p ≤ 0.001 compared to vehicle control using Student’s t-test.

**Table 1.** Collated dissociation constants (K_D_) of the indicated constructs for human FAP, mouse FAP, or human DPP-IV, as determined by biolayer interferometry.

|  | hFAP | mFAP | hDPP-IV |
| --- | --- | --- | --- |
| H4-Fc | 1.41E-11 ± 2.7E-12 | 1.42E-08 ± 3.3E-10 | NA |
| H15-Fc | 1.28E-09 ± 2.89E-11 | NA | NA |
| H17-Fc | 5.39E-10 ± 6.2E-12 | NA | NA |
| NGS2405-Fc | 3.56E-10 ± 8.3E-12 | 7.95E-08 ± 1.1E-08 | NA |

Flow cytometry was used to determine the ability of the lead VNAR-Fc constructs to engage FAP in its native membrane-bound conformation. Towards this aim, we employed CWR-R1^FAP^ cells, a castration-resistant prostate cancer cell line with stable heterologous FAP expression, as well as parental FAP-null CWR-R1 cells. Additionally, we used an immortalized CAF cell line with endogenous FAP expression, hPrCSC-44, and FAP-null PC-3 prostate adenocarcinoma cells. Each of the prostate cancer cell lines were incubated with serially diluted VNAR-Fc constructs, prior to staining with anti-IgG1-phycoerythrin (PE). Compared to unstained control samples, all VNAR-Fc constructs effected a pronounced increase in brightness of PE-stained cell populations when testing CWR-R1^FAP^ and hPrCSC-44 cells, but not in CWR-R1 or PC-3 cells. Cellular binding was both dose-dependent and saturable **(Figure 4H-I, Supplemental Figure 5)**. In this cellular context, all VNAR-Fcs demonstrated specific and high-affinity binding to FAP-positive cells as seen in CWR-R1^FAP^ and hPrCSC-44 cell lines. Negligible binding was observed in FAP-null cell lines, CWR-R1 and PC-3, highlighting the selective recognition of the VNAR-Fc constructs.

### VNAR-Fc constructs internalize in a dynamin-dependent manner

To explore whether the VNAR-Fc constructs were suitable for the development of ADCs, we conducted a series of internalization studies. Since many current ADCs exploit intracellular factors to initiate conditional delivery of cytotoxic drugs, selective internalization into targeted cells is a prerequisite for delivering payloads while sparing healthy tissues. Confocal microscopy studies were performed in the FAP-positive hPrCSC-44 and FAP-null PC-3 cells. Cells were co-incubated with Alexa Fluor 647-labeled VNAR-Fc (VNAR-Fc-AF647, 10nM) and fluorescein-dextran as a marker of fluid-phase endocytic cargo. After a 1-hour incubation, cells were fixed and labeled with membrane and nuclear stains. In hPrCSC-44 cells, each of the lead VNAR-Fc-AF647s displayed intense fluorescent puncta throughout the cytoplasm **(Figure 5A, C, E, G)**. Colocalization of VNAR-Fc-AF647 with endosomal compartments is illustrated using line scans of AF647 and fluorescein fluorescence intensity in multichannel composite images. In contrast, VNAR-Fc-AF647 fluorescence was not detected in PC-3 cells, despite being viable and endocytosis-competent, as indicated by endocytic uptake of fluorescein-dextran **(Supplemental Fig 6)**.

**Figure 5.**
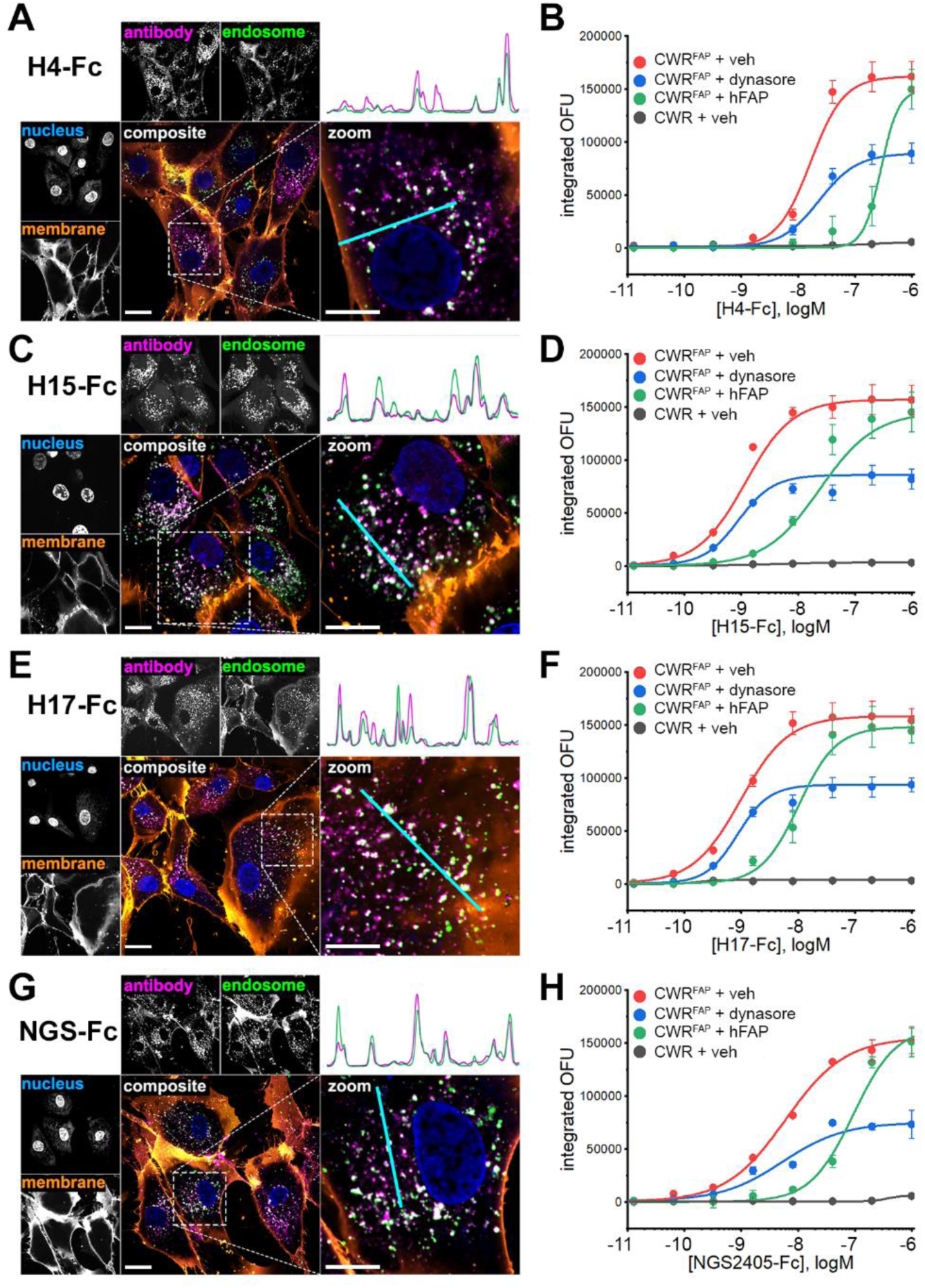
Anti-FAP VNAR-Fc constructs internalize into FAP-expressing cells. **A,C,E,G,** confocal microscopy images of hPrCSC-44 cells after incubation with H4-Fc-AF647 (A), H15-Fc-AF647 (C), H17-Fc-AF647 (E) or NGS2405-Fc-AF647 (G) for 1hr, using 10nM of anti-FAP VNAR-Fc-AF647 and 50µg/ml of fluorescein-dextran. Single-channel images of VNAR-Fc-AF647 localization, fluorescein-labeled endosomes, Hoescht 33342-labeled nuclei, and CellBrite 555-labeled membranes are shown. Merged composite images depicting whole-cells and enlarged regions of interest are shown as colored fluorescence overlays. *Top right*, plots of relative fluorescent signal detected in line scans (teal) in the antibody channel and the endosome channel are shown to illustrate spatial co-localization of punctate structures. Scale bar represents 20µm in uncropped images, and 10µm in zoomed insets. **B, D, F, H,** aggregate data from high-content live-cell imaging of anti-FAP VNAR-Fc internalization into CWR-R1^FAP^ or CWR-R1 cells. Antibodies were directly labeled with pHrodoRed, integrated pHrodoRed fluorescence detected after treatment with the indicated concentration of H4-Fc-pHrodoRed (B), H15-Fc-pHrodoRed (D), H17-Fc-pHrodoRed (F), or NGS2405-Fc-pHrodoRed (H) in CWR-R1 cells or CWR-R1^FAP^ cells that were either treated with DMSO vehicle (0.1%), dynasore (30µM), or 100nM of soluble recombinant hFAP. Data represents mean ± s.e.m. from n=3 independent experiments.

As an orthogonal method of measuring antibody internalization, we performed high content live-cell imaging experiments using VNAR-Fcs directly labeled with the pH-sensitive fluorophore, pHrodoRed. This approach uses pHrodoRed fluorescence to report antibody translocation through the acidic endolysosomal system. VNAR-Fc-pHrodoRed internalization was assessed in FAP-positive CWR-R1^FAP^ cells and in parental FAP-null CWR-R1 cells. In congruence with the confocal studies, each of the four lead VNAR-Fc-pHrodoRed constructs readily internalized into CWR-R1^FAP^ cells **(Figure 5B, D, F, H)**. Fluorescent signal from pHrodoRed labeled antibodies was dose-dependent and saturable, with low nanomolar EC_50_ values **(Table 2)**. Internalization experiments performed in the presence of soluble recombinant hFAP (100nM) resulted in a rightward dose-response shift, dynamics that are characteristic of competitive inhibition occurring between association of the VNAR-Fc constructs with either cellularly expressed FAP or soluble FAP. In contrast, treating cells with dynasore (30µM), an inhibitor of dynamin proteins involved in endocytic vesicle scission, suppressed the maximum penetrance of antibody internalization while the potency of internalization for each VNAR-Fc-pHrodo was unaffected, consistent with non-competitive inhibition. No pHrodoRed signal was detected after incubation of the VNAR-Fc constructs with FAP-null CWR-R1. Altogether, these data demonstrated that internalization of each VNAR-Fc was dependent on antibody concentration, cellular FAP expression, and dynamin activity. Competitive inhibition by soluble FAP further underscores the target-specificity of these results.

**Table 2.** Collated half-maximal effective concentrations (EC_50,_ nM) of the indicated constructs for internalization into CWR-R1^FAP^ or CWR-R1 cells after treatment with either DMSO (0.1%), dynasore (30µM), or soluble recombinant hFAP (100nM). EC_50_ values are derived from logistic fitting of mean ± s.e.m. fluorescence values from n=3 independent biological experiments. NA, EC_50_ not available.

| EC<sub>50</sub>, nM | CWR-R1<sup>FAP</sup> |  |  | CWR-R1 |
| --- | --- | --- | --- | --- |
|  | DMSO | Dynasore | Soluble FAP | DMSO |
| H4-Fc | 16.4 ± 7.7 | 22.3 ± 16 | 285 ± 330 | NA |
| H15-Fc | 1.17 ± 0.14 | 0.886 ± 0.13 | 24.1 ± 2.3 | NA |
| H17-Fc | 0.968 ± 0.23 | 0.857 ± 0.21 | 10.1 ± 11 | NA |
| NGS2405-Fc | 6.22 ± 0.52 | 4.45 ± 1.1 | 80.4 ± 8.9 | NA |

### Functional characterization of lead anti-FAP VNAR-Fc constructs as antibody-drug conjugates

Having validated H4-Fc, H15-Fc, H17-Fc and NGS2405-Fc as high affinity anti-FAP antibodies that selectively internalize into FAP-positive cells, we next aimed to test these agents as vectors for cytotoxic drug delivery. Lead anti-FAP VNAR-Fc constructs were conjugated to monomethyl auristatin E (MMAE), a commonly used antimitotic ADC payload. ADCs were designed with a cleavable glycopeptide linker which enables conditional release of MMAE after endocytic internalization and sequential cleavage by two lysosomal enzymes, β-glucoronidase and cathepsin B. Linker-payloads were incorporated into antibody constructs via site-specific modification of Fc glycans, resulting in a drug-antibody-ratio (DAR) of 2. ADCs were evaluated for their ability to selectively kill FAP-positive CWR-R1^FAP^ and hPrCSC-44 cells, while sparing FAP-null CWR-R1 and PC-3 cells, using apoptosis as a marker of cytotoxicity. Cells were incubated with serially diluted VNAR-Fc-MMAEs, and apoptosis was detected by monitoring signal from a fluorogenic caspase 3/7 substrate. Free unconjugated MMAE toxin and a non-targeting IgG-MMAE served as controls. Free MMAE induced apoptosis in all cell lines regardless of FAP expression. Likewise, maximal concentrations (300nM) of IgG-MMAE produced a non-discriminatory induction of apoptosis in all cell lines, but this effect was not potent enough to determine an EC_50_ **(Figure 6)**. In contrast, the VNAR-Fc-MMAEs selectively and potently induced apoptosis in CWR-R1^FAP^ and hPrCSC-44 cells **(Figure 6A&C)** while mimicking the low potency effects seen with IgG-MMAE in CWR-R1 and PC-3 cells **(Figure 6C&D)**. As an additional means of testing ADC efficacy, we conducted cell killing assays using cellular reductive capacity as a readout for metabolic activity and cell viability, recapitulating the same dose-dependent and selective killing of FAP-expressing CWR-R1^FAP^ but not CWR-R1 cancer cells **(Supplemental Fig. 7)**. EC_50_ values for these assays are collated in **Table 3**. These data establish our VNAR-Fc-MMAEs as potent ADCs for the selective killing of FAP-expressing prostate cancer cells.

**Figure 6.**
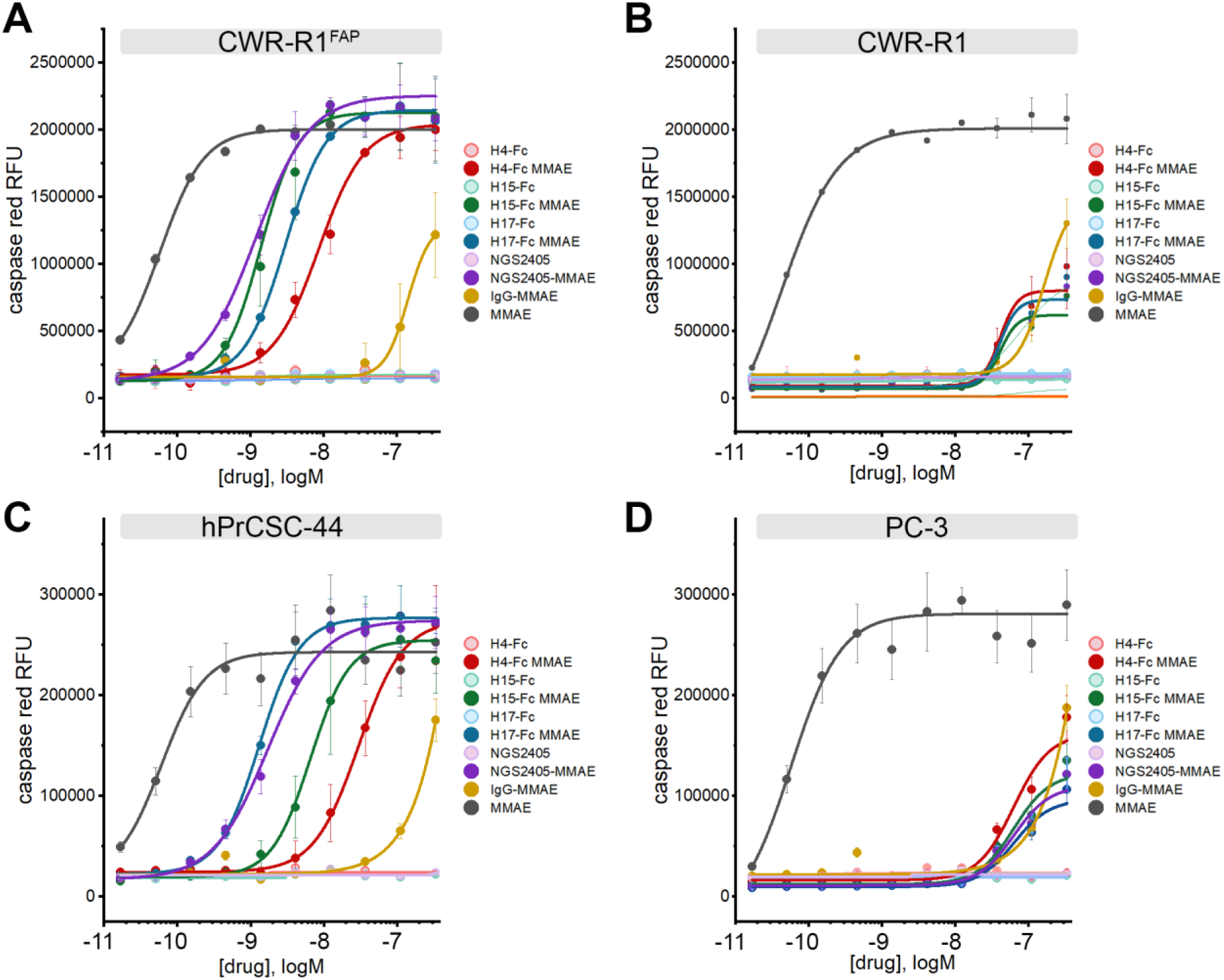
*In vitro* cytotoxicity of anti-FAP VNAR-Fc-MMAE antibody-drug conjugates. Anti-FAP VNAR-Fcs site-specifically conjugated to a monomethyl auristatin E (MMAE) payload were tested for induction of caspase 3/7 activity, as detected using a fluorogenic caspase 3/7 substrate (NucView555) in high-content live-cell imaging experiments. Assays were conducted in parallel with parental unconjugated VNAR-Fc, a non-targeting isotype control VNAR-Fc-MMAE, and free MMAE drug in **A)** FAP-positive CWR-R1^FAP^ cells, **B)** FAP-negative parental CWR-R1 cells, **C)** FAP-positive hPrCSC-44 CAF cells, and **D)** FAP-negative PC-3 cells. Data represents mean ± s.e.m. values from n=3 independent experiments.

**Table 3.**
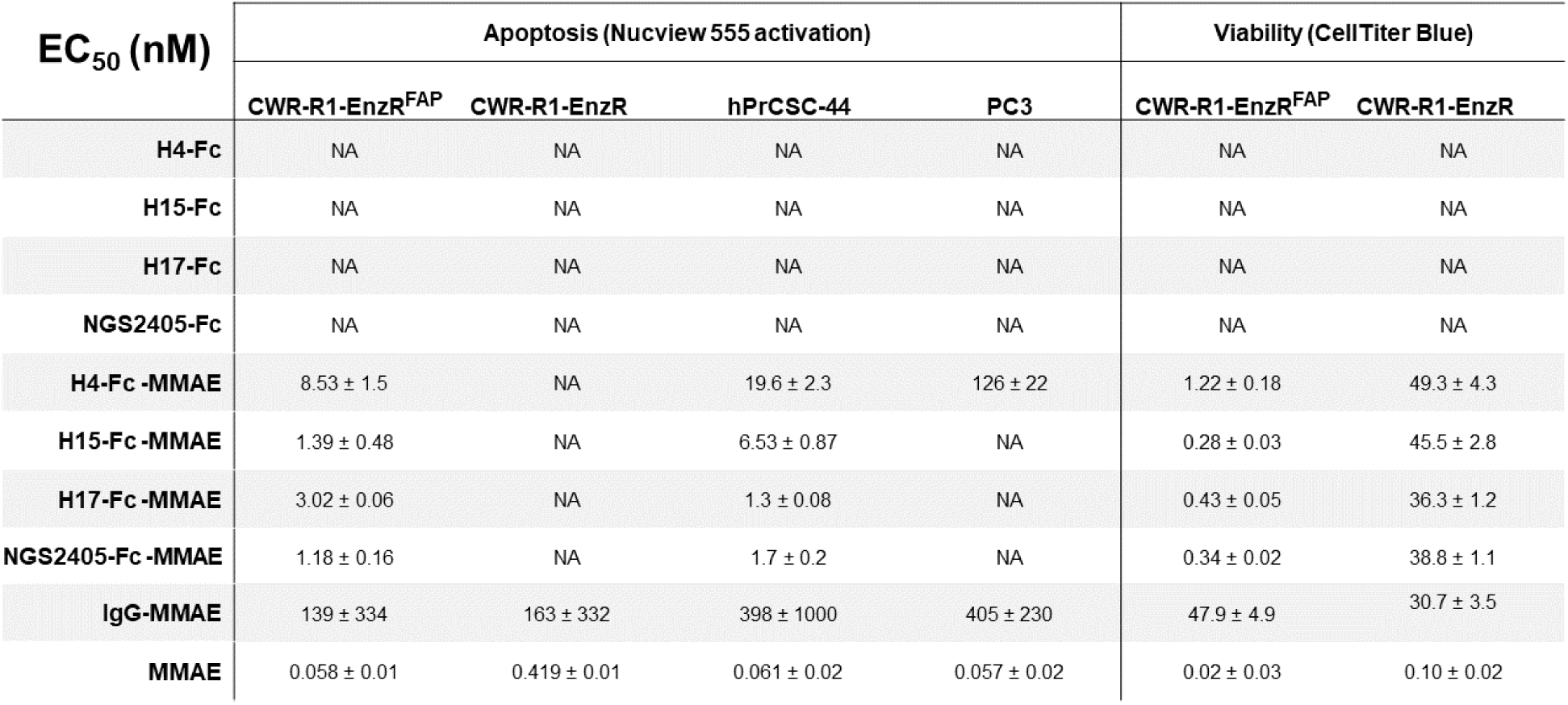
Collated half-maximal effective concentrations (EC_50_, nM) of the indicated constructs for activation of fluorogenic caspase 3/7 activity in the indicated cell lines, as determined by high-content live-cell fluorescence microscopy. EC_50_ values for the induction of cell death, assessed by measuring cellular metabolic activity with CellTiter Blue, are also shown. EC_50_ values are derived from logistic fitting of mean ± s.e.m. absorbance values from n=3 independent biological experiments. NA, EC_50_ not available.

| EC<sub>50</sub> (nM) | Apoptosis (Nucview 555 activation) |  |  |  | Viability (CellTiter Blue) |  |
| --- | --- | --- | --- | --- | --- | --- |
|  | CWR-R1-EnzR<sup>FAP</sup> | CWR-R1-EnzR | hPrCSC-44 | PC3 | CWR-R1-EnzR<sup>FAP</sup> | CWR-R1-EnzR |
| H4-Fc | NA | NA | NA | NA | NA | NA |
| H15-Fc | NA | NA | NA | NA | NA | NA |
| H17-Fc | NA | NA | NA | NA | NA | NA |
| NGS2405-Fc | NA | NA | NA | NA | NA | NA |
| H4-Fc-MMAE | 8.53 ± 1.5 | NA | 19.6 ± 2.3 | 126 ± 22 | 1.22 ± 0.18 | 49.3 ± 4.3 |
| H15-Fc-MMAE | 1.39 ± 0.48 | NA | 6.53 ± 0.87 | NA | 0.28 ± 0.03 | 45.5 ± 2.8 |
| H17-Fc-MMAE | 3.02 ± 0.06 | NA | 1.3 ± 0.08 | NA | 0.43 ± 0.05 | 36.3 ± 1.2 |
| NGS2405-Fc-MMAE | 1.18 ± 0.16 | NA | 1.7 ± 0.2 | NA | 0.34 ± 0.02 | 38.8 ± 1.1 |
| IgG-MMAE | 139 ± 334 | 163 ± 332 | 398 ± 1000 | 405 ± 230 | 47.9 ± 4.9 | 30.7 ± 3.5 |
| MMAE | 0.058 ± 0.01 | 0.419 ± 0.01 | 0.061 ± 0.02 | 0.057 ± 0.02 | 0.02 ± 0.03 | 0.10 ± 0.02 |

### PET/CT imaging of FAP-expressing tumors *in vivo*

The ability for an antibody to selectively localize to targeted tissues *in vivo* is an essential requirement for its utility as a therapeutic or diagnostic agent. To evaluate the potential of our VNAR-Fc constructs as imaging probes, we tested their ability to detect hFAP in a localized *in vivo* model of prostate cancer. Anti-FAP VNAR-Fcs were site-specifically conjugated with a deferoxamine (DFO) chelator and radiolabeled with [^89^Zr] (t_1/2_=3.7 days), enabling PET/CT detection of their biodistribution in treated mice. Mice bearing subcutaneous xenografts composed of either CWR-R1^FAP^ or CWR-R1 prostate cancer cells were injected with either [^89^Zr]Zr-H4-Fc, [^89^Zr]Zr-H15-Fc, [^89^Zr]Zr-H17-Fc, or [^89^Zr]Zr-NGS2405-Fc, and PET/CT scans were serially acquired over 96 hours. Analyses of three-dimensional reconstructions of PET images revealed that each of the imaging probes displayed significant uptake in FAP-expressing tumors compared to control tumors, however nuanced differences in probe pharmacokinetics were discernable **(Figure 7)**. [^89^Zr]Zr-H15-Fc had the most rapid tumor localization, with uptake in FAP-expressing tumors being significantly greater than in negative tumors within 4 hours **(Figure 7D-F)**. In contrast, significant tumor uptake was not measured until the 24-hour time point for the remaining probes. Time-dependent accumulation of [^89^Zr]Zr-H4-Fc, [^89^Zr]Zr-H15-Fc and [^89^Zr]Zr-H17-Fc at FAP-expressing tumor sites was evident over the course of the 96-hour study, while tumor localization of [^89^Zr]Zr-NGS2405-Fc rapidly plateaued within 24-hours. Signal from FAP-expressing tumors in mice injected with [^89^Zr]Zr-H4-Fc or [^89^Zr]Zr-H17-Fc failed to significantly exceed the signal detected in kidneys or livers. In contrast, FAP-expressing tumors were easily resolved using either [^89^Zr]Zr-H15-Fc or [^89^Zr]Zr-NGS2405-Fc, as both probes displayed significantly higher localization to tumor sites compared to all other organs within 24 hours **(Figure 7A-L)**.

**Figure 7.**
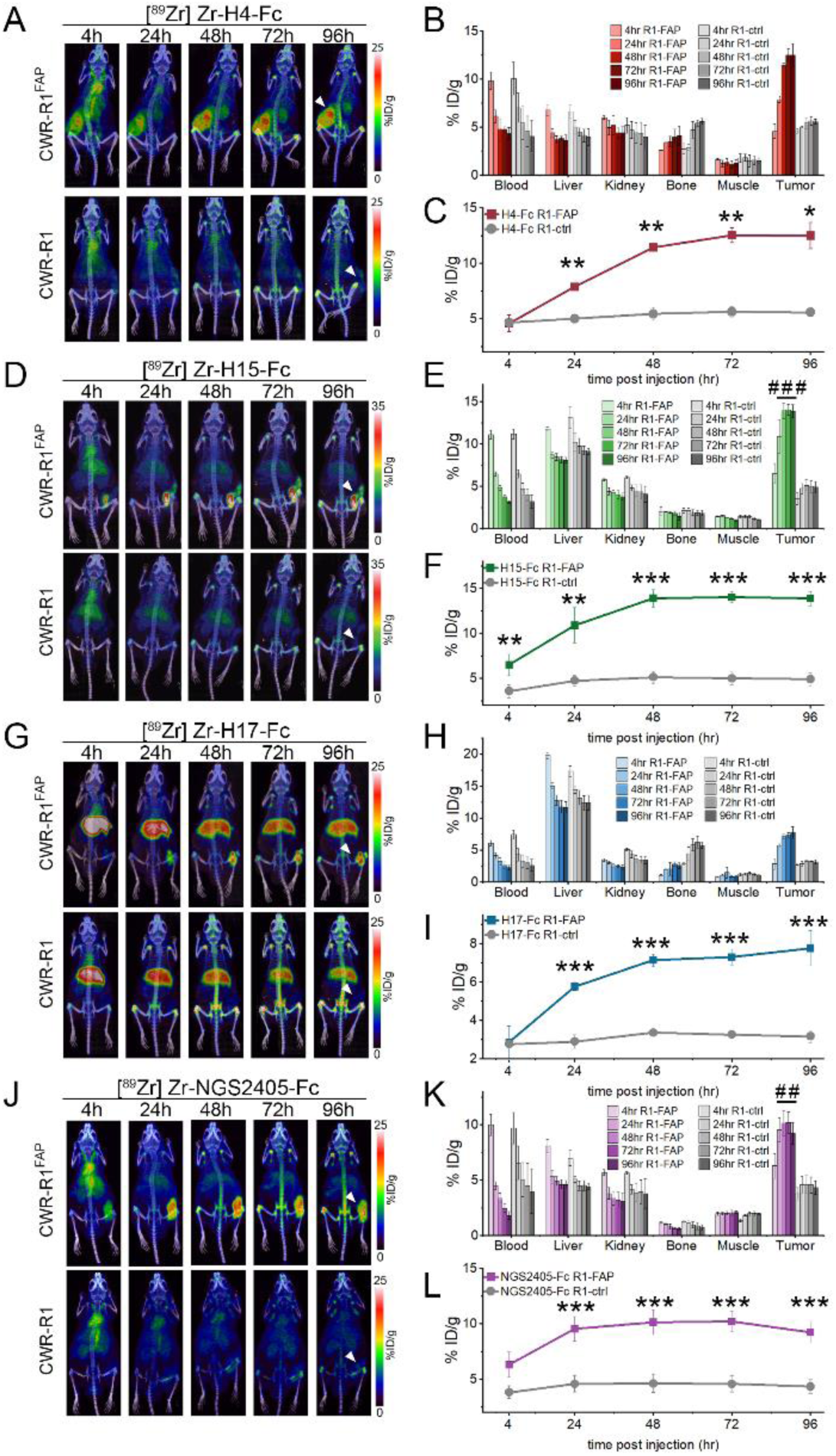
PET/CT imaging of FAP-expressing xenografts *in vivo*. Representative images from PET/CT scans of **A)** [^89^Zr]Zr-H4-Fc, **D)** [^89^Zr]Zr-H15-Fc, **G)** [^89^Zr]Zr-H17-Fc, and **J)** ^89^Zr]Zr-NGS2405-Fc localization in mice bearing CWR-R1^FAP^ (top) or CWR-R1 (bottom) xenografts at the indicated time points. Biodistribution among the indicated organs at the indicated time points in mice bearing either CWR-R1-EnzR^FAP^ or CWR-R1-EnzR prostate cancer xenografts for **B)** [Zr^89^]Zr-H4-Fc, **E)** [Zr^89^]Zr-H15-Fc, **H)** [Zr^89^]Zr-H17-Fc, and **K)** [Zr^89^]Zr-NGS2405-Fc. Quantitative analysis of **C)** [Zr^89^]Zr-H4-Fc, **F)** [Zr^89^]Zr-H15-Fc, **I)** [Zr^89^]Zr-H17-Fc, and **L)** [Zr^89^]Zr-NGS2405-Fc uptake in CWR-R1^FAP^ or CWR-R1 subcutaneous xenografts. Radiolabeled antibodies were delivered via tail vein injection in n=3 mice per condition, *p values, \** p ≤ 0.05; ** p ≤ 0.01, *** p ≤ 0.001 compared to FAP-negative controls. *^##^p* ≤ 0.01; *^###^p* ≤ 0.001 compared to all secondary organs at the same time point.

## Discussion

The TME is a central player in promoting cancer progression and metastasis^34,35^. A greater understanding of the TME and the crosstalk between the individual components will lead to the development of new therapies and the effective implementation of existing immunotherapies. While CAFs exist as heterogenous populations within the TME, CAFs that express FAP display a markedly protumorigenic phenotype^5^. The mechanisms by which FAP-positive CAFs aid cancer progression by promoting immunosuppression, tumor growth, metastasis, and drug-resistance are slowly being elucidated. Recent findings by our group and others suggest their targeted elimination can turn an immunologically cold TME hot by eliciting a proinflammatory response and immune cell infiltration^12,13^. Additionally, FAP-positive CAFs can be leveraged for diagnostic applications. FAP-positive CAFs in the stroma are known to be predictive of poor survival in many cancer types ranging from colon to lung cancer^36–38^. Gallium-68 labelled FAPIs have also been found to be more accurate at detecting disease than the standard-of-care PET probe [^18^F]FDG^39,40^. Imaging FAP in the stroma represents a strategy to overcome tumor heterogeneity since CAFs are more genetically stable and often surround cancer cells that possess heterogeneous antigen expression. Thus, FAP represents a promising theranostic target in solid tumors that has yet to be fully exploited.

Since the publication of the first FAPI more than five years ago, a veritable arms race has occurred resulting in dozens of FAP radioligands with more than 1000 papers published^41^. Only a few studies have been published on the development of biologics targeting FAP^12,42,43^. Several anti-FAP camelid domains have been reported in the literature, but there have been no publications documenting the identification of anti-FAP VNARs from either synthetic or immunized libraries^44,45^. Here, we identified three anti-FAP VNARs, H4, H15, and H17, from a phage display library constructed from the immune cells of a nurse shark immunized with hFAP. Since antibody phage display typically yield titers of >10^5^ clones, this precluded a practical means of screening all putative hits, leading to potential loss of low-frequency binders. For this reason, NGS was used to complement biopanning. From the NGS data of our immunized library, we were able to identify NGS2405, an anti-FAP VNAR with an entirely unique CDR3. As bivalent Fc fusion proteins, the four VNAR-Fc constructs bound hFAP with high affinity while only H4-Fc and NGS2405 were cross-reactive with mouse FAP. The VNAR-Fc constructs also engaged FAP on cell lines engineered to express FAP and immortalized CAF cells. The VNAR-Fc constructs rapidly and efficiently internalized into FAP-expressing cells – a prerequisite for effective theranostics. As a result, the VNAR-Fc constructs were able to eliminate FAP-positive cells *in vitro* as ADCs and detect FAP-positive xenografts *in vivo* by PET imaging.

Our approach for *in silico* VNAR identification was predicated on the idea that immunization would result in clonal expansion of B cells which encode high-affinity anti-FAP IgNARs leading to a concomitant enrichment of functional anti-FAP VNARs in our library^46^. Consistent with this notion, we found a strong correlation between the frequency of FAP-reactive Sanger clones and the abundance of these same clones as detected by NGS of our library . While this observation held true among VNARs with clonotypes that were experimentally identified by phage display, it did not directly translate to enable identification of new anti-FAP VNARs through simple prioritization of highly abundant clones within the NGS dataset. Our initial attempt to identify novel anti-FAP VNARs based solely on NGS clonal frequency failed to identify any binders suggesting that these highly abundant VNARs may be directed against non-specific environmental factors. Using NGS data from an unrelated control VNAR library, we were able to resolve clusters of relatively abundant VNARs that were specifically present in the FAP-immunized library resulting in a vastly improved hit-rate (75%) for NGS-guided antibody discovery. Despite our stringent clonotype cutoff of 85% CDR3 sequence identity, we found log-order differences in affinity among binders within FAP-reactive clonotypes. Our strategy also yielded entirely novel clonotypes in contrast to widely adopted applications of NGS-guided antibody discovery which aim to generate pools of candidate binders based on homology to lead antibodies found during *in vitro* antibody display screening^47–49^. Other recent studies have also identified novel clonotypes from immunized libraries by using NGS to monitor evolution of low-frequency clonotype populations after iterative rounds of affinity-based selection events^50–52^. To our knowledge, our approach is the first method to enable NGS-guided discovery of naturally occurring antibodies without any preliminary rounds of biopanning. Our results are also in broad agreement with reports finding that simple heuristics such as NGS clone abundance are poorly correlated to binding ability when used as a sole metric, but have strong predictive power when coupled with clonotype clustering approaches^53,54^.

We determined that our four VNAR-Fc constructs recognized three separate epitopes through binning experiments. H4-Fc and NGS2405-Fc recognized uniquely independent epitopes on FAP that were conserved across species. While H15-Fc and H17-Fc shared very little CDR3 sequence homology, they still showed overlap to the same conserved epitope on hFAP with high affinity. Interestingly, as PET imaging agents *in vivo*, [^89^Zr]Zr-H15-Fc and [^89^Zr]Zr-H17-Fc behaved quite differently. Tumor uptake for [^89^Zr]Zr-H15-Fc was rapid starting at 4h and persisted out to 96h reaching a final value of 15%ID/g by ROI analysis. The tumor uptake of [^89^Zr]Zr-H17-Fc was slower and reached a %ID/g nearly half that of [^89^Zr]Zr-H15-Fc at 96h. High liver uptake was also observed in the FAP-positive and null xenografts administered [^89^Zr]Zr-H17-Fc which was not seen with the other VNAR-Fc constructs. These results suggest that several factors exist that determine the ability of a VNAR-Fc to localize to a tumor *in vivo*, not just their affinity for the epitope. It is known that the isoelectric point (pI) of an antibody can affect the pharmacokinetics, with high pI antibodies (>8) rapidly localizing to the liver as was observed with [^89^Zr]Zr-H17-Fc^55^. Given the disparity in pI values for H15-Fc and H17-Fc (pI – 6.5 and 8.39, respectively), this is the most li kely explanation for the performance of [^89^Zr]Zr-H17-Fc *in vivo*. The properties of H4-Fc further underscore that tumor localization *in vivo* is multi-factorial. While H4-Fc had the best affinity for FAP of all the constructs (14pM), it had the worst EC_50_ values for internalization and cytotoxicity, yet it demonstrated high tumor uptake. The on-site specificity of H4-Fc and NGS2405-Fc is also notable, as both antibodies are cross-reactive with endogenous mFAP, yet displayed low secondary accumulation as PET agents, enabling easy resolution of FAP-positive tumors. This further documents that FAP-positive cells are absent in normal in tissues and organs and that the xenograft model we used had limited stroma with few FAP-positive murine CAFs as previously described^42^.

Previous attempts targeting FAP for therapeutic benefit centered on inhibiting the proteolytic activity of FAP with the peptidyl boronic acid inhibitor Talabostat. Though preclinical studies in animal models were promising, Talabostat met with little success in the clinic documenting that the inhibition of FAP alone was not an effective strategy^56^. We found that none of the VNAR-Fc constructs inhibited the enzymatic activity of FAP. The epitopes recognized by our VNARs did not occlude the substrate specificity pocket or catalytic triad. Rather than inhibiting the activity of FAP, our strategy was to use FAP to gain entry into CAFs and deposit cytotoxins with our internalizing VNAR-Fc constrcuts. When armed with the microtubule inhibitor MMAE at a DAR of 2, the VNAR-Fc constructs were effective ADCs eliminating FAP-positive cells with pM to low nM EC_50_ values. All the VNAR-Fc-MMAE ADCs were more cytotoxic by the Cell Titer Blue assay than our previously published humanized anti-FAP ADC huB12-MMAE with a DAR of 4^12^. HuB12 internalized less efficiently (EC_50_ - 17.3nM) than the VNAR-Fc constructs potentially explaining this difference. Our approach differs from previous strategies using biologics targeting FAP. The first biologic evaluated in the clinic for FAP was the humanized antibody sibrotuzumab. Sibrotuzumab proved safe and well tolerated in phase I trials, however lacked significant efficacy in phase II trials^23,57^. *In vitro* studies revealed poor internalization kinetics of sibrotuzumab, a finding that could potentially explain its poor clinical efficacy. In the past decade, Roche developed several FAP targeted biologics including the engineered IL2 - FAP antibody fusion protein simlukafusp alfa and several bispecifics that bind DR5 (RG7386), CD40 (RG6189), and 4-1BB (RG7827) in addition to FAP^25,58^. Two of the bispecifics, RG7386 and RG6189, have been discontinued while RG7827 is undergoing a phase I trial. It is also debatable whether trying to stimulate the immune system with bispecific antibodies in a highly immunosuppressive TME possessing a few immune cells is the most prudent strategy.

VNARs occupy a unique niche in biologics and are markedly different from other binding domains including camelids. Lacking a CDR2, VNARs have two additional hypervariable loops (HV2 and HV4) resulting in four loops of diversity in their compact structure^59^. Our NGS data shows that CDR3 lengths of up to 25 amino acids or more are also common with VNARs. Through intricate disulfide bonding patterns, VNARs can adopt complex geometries, providing a molecular dexterity that is absent in other antibody domains^60^. Because of their small size, modular design, and high solubility, VNARs can be easily engineered to create hypervalent and multi-paratopic targeting vectors that would be near impossible to create using conventional antibody domains. Additionally, studies in rodent models, and a recent phase I clinical trial with a VNAR-derived therapeutic for lung fibrosis (NCT05914909), have documented that VNARs are non-immunogenic with no significant generation of anti-drug antibodies in rodents or humans^61–63^. Coupled with their ease of production, scalability, and cost-effectiveness, VNAR-Fc constructs represent a readily translatable biologic with the potential to dramatically alter how antigens are targeted in the future.

## Acknowledgements

This project was supported in part by NIH/NCI R01 CA237272 (A.M. LeBeau), NIH/NCI R01 CA233562 (A.M. LeBeau), a 2018 Prostate Cancer Foundation Challenge Award (A.M. LeBeau), a Prostate Cancer Foundation Young Investigator Award (A.M. LeBeau), Andy North and Friends (A.M. LeBeau), the Randy Shaver Cancer Research and Community Fund (A.M. LeBeau), and the Department of Veterans Affairs, Veterans Health Administration, Office of Research and Development, Clinical Sciences Research and Development Merit Award CX002479 (D. Kosoff).

## Materials and Methods

### Shark housing

Sharks were housed in a fiberglass round tank with an operational volume of 18m^3^, and was operated as a recirculating aquaculture system with water turnover occurring approximately twice per hour. Water salinity (32-35 ppt), temperature (27° C), ammonia content (<0.03 ppm), nitrogen dioxide (<0.1 ppm) nitrate (<0.1 ppm), pH (7.8-8.3), and dissolved oxygen (>5.0mg/l) were monitored daily. Sharks were monitored weekly by veterinary staff for signs of illness or parasitism, and were fed 3 times per week on a diet consisting of octopus, sea bass, shrimp and mackerel at a rate of 4% body weight per week.

### Immunization

Sharks weighing between 3-4kg were sedated using 0.01% MS-222 (Syndel, Ferndale, WA) solubilized in artificial seawater. Blood was drawn from the caudal vein using a syringe pre-filled with 1ml of 1% EDTA and fitted with a 20G needle. Blood was collected two weeks prior to beginning immunization programs (pre-bleed) and at two-week intervals during immunization for 10 weeks. Initial immunizations were performed using 250µg antigen emulsified 1:1 with Complete Freund’s Adjuvant (CFA), which was administered subcutaneously in the lateral fin. At two-week intervals, sharks were boosted with a tapering dose of antigen (250µg - 50µg). Antigen was emulsified in Incomplete Freund’s Adjuvant (IFA) at a 1:1 ratio and delivered subcutaneously into alternating lateral fins on weeks 2, 6 and 8. On week 4, purified antigen suspended in PBS supplemented with 350 mM urea was delivered intravenously into the caudal vein. Plasma and buffy coat fractions were isolated from samples of whole blood after centrifugal fractionation at 300xRCF for 5 min at 4°C with rotor brakes turned off. Immunization was monitored by screening plasma samples for the presence of convalescent IgNARs that are immuno-reactive to the antigen of interest, using biolayer interferometry.

### VNAR phage-display library construction

Shark plasma samples that produced a strong anti-FAP BLI response (response > 0.5nm) were used to identify corresponding buffy coat samples from the same time points, which were used for library assembly. RNA was isolated from buffy coat samples using an RNAeasy kit (Qiagen, Hilden, Germany), and converted to cDNA using a High Capacity RNA-to-cDNA kit (Applied Biosystems, Thermo Fisher Scientific, Waltham, MA). Custom oligonucleotide primers specific to the framework 1 and framework 4 regions of IgNAR V regions were used in polymerase chain reaction (PCR) experiments to specifically amplify VNAR-encoding cDNA. PCR amplicons were separated by agarose gel electrophoresis, nucleic acid bands of approximately 350bp were excised, purified, restriction digested, and ligated into a pADL-22c phagemid vector (Antibody Design Labs, San Diego, CA) using standard methods. PCR, cDNA synthesis, and ligation steps were each performed in 48 independent reactions. Groups of 4 ligation reactions were pooled, generating 12 pools of VNAR-encoding phagemids, which were each used in independent electroporations. Electrocompetent TG-1 cells (Lucigen, Madison, WI) were electroporated, plated on selective media dishes and incubated overnight at 30°C. Library size was titred by serial dilution of pooled transformants. After overnight growth, E. coli bacterial lawns were scraped, pooled, supplemented with 20% glycerol, OD_600_ was measured, and E. coli stocks of the immunized anti-FAP VNAR phagemid library were aliquoted and stored at -80°C. Phage was produced from E. coli stocks of the immunized anti-FAP VNAR phagemid library using standard phage rescue methods and purified by repeated precipitation with 4% (w/v) polyethylene glycol 8000 and 0.5M NaCl. Purified phage was titred by serial dilution, infection of naïve TG-1 E. coli cultures, and plating on selective media plates.

### Phage display biopanning

Phage purified from an immunized anti-FAP VNAR phage display library was used to identify clones against recombinant human FAP. Recombinant human FAP (FAP-H5244, Acro Biosystems, Newark, DE) was biotinylated using EZ-link NHS-PEG4-biotin (Thermo Fisher Scientific, Waltham, Mass) according to the vendor recommendations. A volume of purified phage representing 1000x fold more particles than the E. coli library size was diluted in PBS to a volume of 1 ml. Phage with non-specific affinity for bead and tube materials were precleared by end-over-end incubation with Dynabeads M-270 Streptavidin (Invitrogen, Carlsbad, CA) in microcentrifuge tubes. Soluble phage was transferred to a new tube containing soluble biotinylated recombinant FAP, supplemented with 2% (w/v) BSA. Phage and biotinylated FAP were incubated for 1hr with end-over-end mixing, phage/FAP complexes were transferred to a new tube containing Dynabeads M-270 streptavidin, and complexes were incubated for an additional 30min with end-over-end mixing. The remaining biopanning protocol was performed using standard methods as previously described by others^64^. Positive binders to recombinant human FAP were readily identified by ELISA screening after a single round of biopanning.

### ELISA

VNARs were produced for 192 individual clones using 5 mM IPTG induction in microtiter plates. Soluble hemagglutinin (HA)-tagged VNARs that leaked into culture media supernatant were screened for binding to hFAP by ELISA. MaxiSorp plates (Nunc Cell Culture, Thermo Fisher Scientific, Waltham, MA) were coated with 50ul of streptavidin (50 µg/ml in PBS, Promega, Madison, WI) overnight at 4° C. Wells were washed two times with PBS and blocked with 370 μL of 2% BSA in PBS for 1 hour at room temperature. The wells were washed three times with PBS, 0.005% Tween20. 50 μL of biotinylated FAP (1 μg/mL in PBS, 1% BSA, 0.005% Tween20) was added to each well. Plates were shaken at room temperature for 1 hour. The wells were washed three times with PBS, 0.005% Tween20 and the supernatants of VNAR induced cultures were added to each well and shaken at room temperature for 1 hour. The wells were washed three times with PBS, 0.005% Tween20. VNAR binding was detected with a 1:1000 dilution of anti-HA-tag monoclonal antibody conjugated to peroxidase (12-013-819-001, Sigma-Aldrich, St. Louis, Mo.) in PBS, 1% BSA and 50 μL Turbo™ B reagent (Pierce Protein Biology, Thermo Fisher Scientific, Waltham, Mass.). Reactions were stopped with 10 μL of 2.5 M H_2_SO_4_ and the absorbance was measured at 450 nm using a microplate reader. Confirmed positive clones for FAP were sequenced to identify unique clones.

### Next Generation Sequencing

VNAR-encoding DNA was digested out of pADL-22c phagemid vector using SfiI restriction enzyme (New England Biolabs). Excised DNA was purified by agarose gel electrophoresis. VNAR sequences were ligated to Illumina adaptor and sequenced using Illumina MiSeq 2 x 300 bp paired. FASTQ data was trimmed with Skewer to remove adapters sequences and low-quality bases. Paired-end data was merged with FLASH. Data was filtered, keeping data that were flanked by conserved 5’-GCCATGGCTGCTCGAGTGGACCAAACACCGCGTGACTGTGAATGGCCCGGGAGGCCA - 3’ sequences. Nucleotides were translated to amino acid sequences. Sequences with internal stop codons were filtered out. VNAR subtypes were classified based on positioning of cysteine residues, with type‘We trimmed the fastq data with skewer to remove adapters and low-quality bases. We merged the paired-end data with flash. We then filtered the sequences to keep reads that had both of the expected fragment ends and could be translated in frame between expected fragment ends. We translated the nt sequences to protein and then use the position of cysteine residues to subdivide the output into categories. We took then looked for overlap between the two libraries.

### Production of VNAR-Fc antibodies

DNA sequences encoding lead anti-FAP VNAR-Fcs were codon optimized for expression in Chinese hamster (*Cricetulus* griseus) systems and synthesized as double stranded DNA gBlocks (Integrated DNA Technologies, Coralville, IO), prior to integration into a mammalian expression vector encoding the Fc domain of human IgG1 (TGEX-SCblue, Antibody Design Labs) by homologous recombination, using an In-Fusion HD cloning kit (Takara Bio USA, San Jose, CA). Plasmid constructs encoding VNAR-Fcs were transfected into ExpiCHO-S cells using an Expifectamine CHO Transfection Kit (Gibco, Thermo Fisher Scientific, Waltham, MA). Transfection and ExpiCHO-S cell culture maintenance was performed as recommended by the manufacturer. 12-14 days post transfection, cultures were harvested and centrifuged at 2000xRCF for 10 min at 4 °C. Protein-containing supernatant was collected and clarified by centrifugation at 20,000xRCF for 30 min at 4 °C, supplemented with 300mM NaCl and 20mM Na_2_PO_4,_ pH was adjusted to 6.8, and solutions were passed through a 0.22 µm sterile filter. Antibodies were captured using protein A affinity chromatography and further purified by size exclusion chromatography. MabSelect PrismA columns (Cytiva, Marlborough, MA) were equilibrated with five column volumes (CVs) of PBS. Clarified ExpiCHO-S supernatant was onto the protein A columns at 1 CV/min. Unbound protein was washed from the column using 10 CVs of PBS supplemented with 350 mM NaCl. Antibodies were eluted in 7 CVs of 200 mM glycine at pH 3.0, followed by 2.5 CVs of PBS. Eluate was immediately neutralized using 2 M Tris HCl at pH 8.6. Eluates were concentrated and buffer exchanged into PBS using an Amicon stirred chamber with a 30 kDa MWCO ultrafiltration membrane (Millipore). Size exclusion chromatography was performed on an ÄKTApure fast protein liquid chromatography system using a HiLoad 16/600 Superdex 200 pg (Cytiva) column equilibrated with PBS. Samples were loaded and fractionated using a mobile phase of PBS at 0.5 mL/min; chromatograms were obtained by monitoring UV absorbance at 280 nm. Eluted protein fractions contributing to a single peak of UV absorbance corresponding to the theoretical molecular mass of each VNAR-Fc were collected and pooled. Eluates were diluted to 1 mg/mL in PBS, dispensed into 0.5 mL aliquots, flash frozen and stored at -80°C.

### Biolayer Interferometry

All BLI studies were performed in an assay buffer consisting of PBS supplemented with 1% BSA. Recombinant human FAP (FAP-H5244, Acro Biosystems), mouse FAP (FAP-M53H3, Acro Biosystems), and human DPP-IV (DP4-H5221, Acro Biosystems) were each biotinylated using EZ-link NHS-PEG4-biotin (Thermo Fisher Scientific) according to the vendor recommendations. *Screening shark plasma samples*, biotinylated human FAP protein was immobilized on Octet streptavidin (SAX) biosensors (Sartorious, Göttingen, Germany), and exposed to plasma samples from various time points diluted 1:200 in assay buffer. Convalescent IgNARs were allowed to associate with immobilized FAP for 30 min, samples that maintained a signal > 0.5 nm after 30 min of dissociation in assay buffer were determined to be indicative of an anti-FAP immune response. *Determination of purified VNAR-Fc dissociation constants*, hydrated SAX sensors were equilibrated for 60 sec in assay buffer before loading of biotinylated bait protein (30 nM) for 90 sec, followed by a baseline equilibration for 60 sec in assay buffer. Association of serially diluted VNAR-Fc proteins to biosensors with immobilized bait protein was monitored for 5-20 minutes, dissociation of VNAR-Fcs in assay buffer was monitored for an equivalent period of time. Prior to each experiment, unloaded SAX sensors were exposed to 1 µM purified VNAR-Fc to verify lack of binding to empty sensors. Each experiment included a reference well with no analyte, to ensure specificity of signal on the bait-loaded sensors. Binding affinities were determined by kinetic analysis of binding curves in the Octet Data Analysis software (v12.0.2.3), using conditions reflecting bivalent analyte and background subtracting data from reference wells. *Antibody cross-competition epitope binning,* hydrated SAX biosensors were monitored during a 60 sec baseline in assay buffer, loaded with biotinylated human FAP (30 nM, 90 sec), followed by exposure to a saturating concentration of a single primary VNAR-Fc (1 µM) for 4 min. A second baseline in assay buffer was recorded for 30 sec, prior to a secondary association step in which biosensors were exposed to a saturating concentration of each individual VNAR-Fc (1 µM). Assays were separately performed using each lead VNAR-Fc as the primary binder. Compared to the average signal recorded during the second baseline step, antibodies that produced a peak BLI signal < 0.1 nm during the secondary association were determined compete with the primary antibody for the same epitope.

### Flow Cytometry

For flow cytometry studies, CWR-R1, CWR-R1^FAP^, hPrCSC-44, and PC3 cell lines were harvested and suspended in flow cytometry staining buffer (eBioscience, Invitrogen) at a final concentration of 1 x 10^6^ cells/mL. Cells were then incubated with various concentrations of VNAR-Fcs for 1 hour on ice. After incubation, cells were centrifuged at 400*g* for 4 minutes and washed with staining buffer thrice. Pelleted cells were then resuspended in staining buffer containing PE labeled goat anti-human IgG Fc (5 μg/mL) (eBioscience, Invitrogen) for 45 min on ice. Cells were washed and resuspended in staining buffer thrice. Samples were analyzed on a Attune Flow Cytometer (ThermoFischer) and fluorescence intensity analysis and histograms generation were done using FlowJo software (Tree Star, Inc.).

### Antibody Internalization Studies

Antibodies were fluorescently labeled using succinimidyl esters of Alexa Fluor 647 (Thermo Fisher Scientific #A20006) or pHrodo Red (P36600) prior to analysis by confocal microscopy or Incucyte imaging, respectively. For confocal microscopy studies, hPrCSC-44 and PC3 cells were seeded onto glass-bottom 35 mm dishes (MatTek, P35GCOL-0-14-C) at a density of 15,000 cells per dish, 48 hours prior to imaging. On the day of analysis, cells were coincubated with Alexa Fluor 647-labeled antibody (10 nM) and fluorescein-labeled dextran (50 µg/mL; Thermo Fisher Scientific #D1821) for 1 hour at 37°C. Cells were then washed three times in PBS and fixed in 4% paraformaldehyde/PBS for 10 minutes. After fixation, cells were stained for 20 minutes with CellBrite Orange (Biotium #30022) and 2 µmol/L Hoescht 33342 (Thermo Fisher Scientific H3570), in accordance with the vendor recommendations. Samples were mounted on a Nikon Eclipse Ti2 inverted microscope equipped with a Yokagawa W1 CSU spinning disk, cells were imaged with a Plan-Apochromat 60x/1.42 oil objective, fluorescence was recorded using a Hamamatsu ORCA-Quest qCMOS camera. Image processing was done in ImageJ 1.54d. For Incucyte internalization experiments, CWR-R1 and CWR-R1^FAP^ cells were suspended in growth medium supplemented with 25 mmol/L HEPES buffer (pH 7.4) and seeded into optical grade 96-well plates (Thermo Fisher Scientific #165305) at a density of 25,000 cells per well. The following day, cells were treated with 0.1% DMSO, 30 µM dynasore (Tocris Bioscience, 2897), or recombinant soluble FAP (100 nM final concentration). Serially diluted pHrodoRed-labeled antibodies were added to cells, pHrodoRed fluorescence was monitored over 3 days in an Incucyte SX5 (Sartorius) using a 20x phase contrast objective and an orange fluorescence optical module (λ_ex_ 557 ± 11nm, λ_em_ 607.5 ± 31.5nm). Data were analyzed by using the Incucyte SX5 Adherent Cell-by-Cell analysis module to measure total integrated orange fluorescence intensity in each experimental condition, values were plotted in OriginLab 2023b for curve fitting.

### Antibody-drug conjugate cytotoxicity assays

ADCs were designed with a glycopeptide cleavable linker requiring cleavage by β-glucuronidase and cathepsin B to enable release of cytotoxic monomethyl auristatin E (MMAE) payloads. ADCs were generated using purified VNAR-Fc antibodies, and were site-specifically conjugated to MMAE using a GlyClick ADC kit (Genovis) according to the manufacturers protocol. Assessment of anti-FAP ADC-induced apoptosis in prostate cancer cell lines was performed using FAP-positive CWR-R1^FAP^ and hPrCSC-44 cells, as well as FAP-negative CWR-R1 and PC3 cells as comparative controls. Cells were suspended in growth medium and seeded into optical grade 96-well plates (Thermo Fisher Scientific #165305) at a density of 25,000 cells per well. The following day, cells were treated with serially diluted unconjugated anti-FAP VNAR-Fc, ADC VNAR-Fc-MMAE, non-targeting control IgG-MMAE, or MMAE. Wells were supplemented with a fluorogenic caspase 3/7 substrate, NucView 530 (Biotium, 10406). NucView 530 fluorescence was monitored over 3 days in an Incucyte SX5 (Sartorius) using a 20x phase contrast objective and an orange fluorescence optical module (λ_ex_ 557 ± 11nm, λ_em_ 607.5 ± 31.5nm). Data were analyzed by using the Incucyte SX5 Adherent Cell-by-Cell analysis module to measure total integrated orange fluorescence intensity in each experimental condition. To measure viability of cells using a metabolic readout, cells were treated with ADC as described above, and were incubated for 72 hours prior to measuring the reductive capacity of cells in each condition using a CellTiter-Blue kit (Promega, G8080), as described in the vendor protocol. Experimental data were plotted in OriginLab 2023b for curve fitting.

### Animal Models

All animal studies were approved by the University of Wisconsin Institutional Animal Care and Use Committee. All animal studies were performed in 3 to 4 weeks old Athymic Nude-Foxn1nu mice (Envigo). Animals (*n* = 3 per experimental group) received subcutaneous injections of either CWR-R1 or CWR-R1^FAP^ cells (1 × 10^6^ cells in 100 µL) suspended in a 1:1 mixture of PBS and Matrigel (Corning). The cell-matrigel mixture was injected into the rear flank of the mice using a 26-gauge needle. Tumor volumes were measured twice weekly with calipers and tumors were allowed to grow to a size of 100–300 mm^3^ before nuclear imaging experiments.

### Bioconjugation and radiochemistry

For nuclear imaging studies, VNAR-Fc antibodies were site-specifically conjugated to deferoxamine (DFO) using a GlyClick DFO kit (Genovis) according to the manufacturers protocol. Zirconium-89 [^89^Zr] used for radiolabeling was provided by the University of Wisconsin Medical Physics Department (Madison, WI). [^89^Zr] Zr-oxalate in 1.0 mol/L oxalic acid was adjusted to pH 7.5 with 2.0 mol/L HEPES. To radiolabel the VNAR-Fcs, the VNAR-Fc-DFO conjugate in PBS (pH 7.5) was added to neutralized [^89^Zr] Zr-oxalate solution (80 μg per 37 MBq) and incubated at 32 °C, shaking at 250 RPM for 1 hour. The labeled product was purified using a size-exclusion PD-10 column preequilibrated with PBS buffer.

### PET/CT image acquisition and analysis

All microPET/CT studies were performed on an Inveon uPET/CT Scanner (Siemens Medical Solutions). Mice (n=3 for each experimental group) were intravenously injected with approximately 4.2 MBq–6.4 MBq of radiolabeled VNAR-Fcs ([^89^Zr]Zr-H4-Fc, [^89^Zr]Zr-H15-Fc, [^89^Zr]Zr-H17-Fc, [^89^Zr]Zr-NSG2405-Fc) for PET studies. PET list mode data were acquired at various time points post injection (4, 24, 48, 72, and 96 h) for 80 million counts using a gamma ray energy window of 350-650 KeV and a coincidence timing window of 3.438ns. A CT-based attenuation correction was performed for approximately 10 minutes with 80 kV, 1mA, 220 rotation degrees in 120 rotation steps, 250ms exposure time, and subsequently reconstructed using a Shepp-Logan filter with 210 micron isotropic voxels. Scans were reconstructed using 3-dimensional ordered-subset expectation maximization (2 iterations, 16 subsets) with a maximum a posteriori probability algorithm (OSEM3DMAP). Two-dimension (2D) images and maximum intensity projections (MIPs) were prepared in Inveon Research Workplace. Quantitative region of interest (ROI) analysis of the PET images was performed using the Inveon Research Workstation software, with values reported in percent injected dose per gram of tissue (% ID/g). Time activity curves were constructed based on the quantification of volumes of interest.

### Cell Culture

All cancer cell lines used in this study were purchased from American Type Culture Collection (ATCC) and were maintained in their respective recommended media, supplemented with 10% FBS (Gibco Thermo Fisher Scientific, Waltham, MA), 1% antibiotic-antimycotic (Gibco), and 1% glutaMAX (Gibco) at 37° C. and 5% CO_2_, unless otherwise specified. Engineered CWR-R1^FAP^ cells were generated and cultured as previously described^42^. FAP-positive hPrCSC-44 cancer associated fibroblast cells were obtained from Dr. W.N. Brennen and have been previously described^65^. hPrCSC-44 cells were maintained in Rooster High Performance Media (Rooster Bio, Frederick, MD) supplemented with 10% Rooster Booster (Rooster Bio). All cell lines have been authenticated using short-tandem repeat profiling, and are routinely monitored for mycoplasma contamination.

## Notes

### Competing Interest Statement

GSG, JPG, and AML are listed as inventors on a pending patent regarding this work

## References

1. Tormoen, G.W., Crittenden, M.R. & Gough, M.J. Role of the immunosuppressive microenvironment in immunotherapy. Adv Radiat Oncol 3, 520–526 (2018).

2. Tie, Y., Tang, F., Wei, Y.Q. & Wei, X.W. Immunosuppressive cells in cancer: mechanisms and potential therapeutic targets. J Hematol Oncol 15, 61 (2022).

3. Tiwari, A., Trivedi, R. & Lin, S.Y. Tumor microenvironment: barrier or opportunity towards effective cancer therapy. J Biomed Sci 29, 83 (2022).

4. Taylor, R.A. & Risbridger, G.P. Prostatic tumor stroma: a key player in cancer progression. Curr Cancer Drug Targets 8, 490–497 (2008).

5. Glabman, R.A., Choyke, P.L. & Sato, N. Cancer-Associated Fibroblasts: Tumorigenicity and Targeting for Cancer Therapy. Cancers (Basel)14(2022).

6. Nyberg, P., Salo, T. & Kalluri, R. Tumor microenvironment and angiogenesis. Front Biosci 13, 6537–6553 (2008).

7. Chen, X. & Song, E. Turning foes to friends: targeting cancer-associated fibroblasts. Nat Rev Drug Discov 18, 99–115 (2019).

8. Barrett, R.L. & Pure, E. Cancer-associated fibroblasts and their influence on tumor immunity and immunotherapy. Elife 9(2020).

9. Chhabra, Y. & Weeraratna, A.T. Fibroblasts in cancer: Unity in heterogeneity. Cell 186, 1580–1609 (2023).

10. Mao, X., et al. Crosstalk between cancer-associated fibroblasts and immune cells in the tumor microenvironment: new findings and future perspectives. Mol Cancer 20, 131 (2021).

11. Takahashi, H., et al. Cancer-associated fibroblasts promote an immunosuppressive microenvironment through the induction and accumulation of protumoral macrophages. Oncotarget 8, 8633–8647 (2017).

12. Gallant, J.P., et al. Mechanistic Characterization of Cancer-associated Fibroblast Depletion via an Antibody-Drug Conjugate Targeting Fibroblast Activation Protein. Cancer Res Commun 4, 1481–1494 (2024).

13. Akai, M., et al. Fibroblast activation protein-targeted near-infrared photoimmunotherapy depletes immunosuppressive cancer-associated fibroblasts and remodels local tumor immunity. Br J Cancer 130, 1647–1658 (2024).

14. Xin, L., et al. Fibroblast Activation Protein-alpha as a Target in the Bench-to-Bedside Diagnosis and Treatment of Tumors: A Narrative Review. Front Oncol 11, 648187 (2021).

15. Hintz, H.M., et al. Imaging Fibroblast Activation Protein Alpha Improves Diagnosis of Metastatic Prostate Cancer with Positron Emission Tomography. Clin Cancer Res 26, 4882–4891 (2020).

16. Brennen, W.N., Isaacs, J.T. & Denmeade, S.R. Rationale behind targeting fibroblast activation protein-expressing carcinoma-associated fibroblasts as a novel chemotherapeutic strategy. Mol Cancer Ther 11, 257–266 (2012).

17. Huang, R., et al. FAPI-PET/CT in Cancer Imaging: A Potential Novel Molecule of the Century. Front Oncol 12, 854658 (2022).

18. Mori, Y., Kratochwil, C., Haberkorn, U. & Giesel, F.L. Fibroblast Activation Protein Inhibitor Theranostics: Early Clinical Translation. PET Clin 18, 419–428 (2023).

19. Poplawski, S.E., et al. Preclinical Development of PNT6555, a Boronic Acid-Based, Fibroblast Activation Protein-alpha (FAP)-Targeted Radiotheranostic for Imaging and Treatment of FAP-Positive Tumors. J Nucl Med 65, 100–108 (2024).

20. Calais, J. FAP: The Next Billion Dollar Nuclear Theranostics Target? J Nucl Med 61, 163–165 (2020).

21. Kratochwil, C., et al. (68)Ga-FAPI PET/CT: Tracer Uptake in 28 Different Kinds of Cancer. J Nucl Med 60, 801–805 (2019).

22. Pang, Y., et al. PET imaging of fibroblast activation protein in various types of cancers by using (68)Ga-FAP-2286: Comparison with (18)F-FDG and (68)Ga-FAPI-46 in a single-center, prospective study. J Nucl Med (2022).

23. Hofheinz, R.D., et al. Stromal antigen targeting by a humanised monoclonal antibody: an early phase II trial of sibrotuzumab in patients with metastatic colorectal cancer. Onkologie 26, 44–48 (2003).

24. Melero, I., et al. A first-in-human study of the fibroblast activation protein-targeted, 4-1BB agonist RO7122290 in patients with advanced solid tumors. Sci Transl Med 15, eabp9229 (2023).

25. Brunker, P., et al. RG7386, a Novel Tetravalent FAP-DR5 Antibody, Effectively Triggers FAP-Dependent, Avidity-Driven DR5 Hyperclustering and Tumor Cell Apoptosis. Mol Cancer Ther 15, 946–957 (2016).

26. Waldhauer, I., et al. Simlukafusp alfa (FAP-IL2v) immunocytokine is a versatile combination partner for cancer immunotherapy. MAbs 13, 1913791 (2021).

27. Bendell, J., et al. Phase 1 trial of RO6874813, a novel bispecific FAP-DR5 antibody, in patients with solid tumors. Molecular Cancer Therapeutics 17(2018).

28. Chung, V., et al. phase Ib/II, open-label, randomised evaluation of atezolizumab plus RO6874281 vs control in MORPHEUS-pancreatic ductal adenocarcinoma. Annals of Oncology 31, S218–S218 (2020).

29. Reichen, C., et al. FAP-mediated tumor accumulation of a T-cell agonistic FAP/4-1BB DARPin drug candidate analyzed by SPECT/CT and quantitative biodistribution. Cancer Research 78(2018).

30. Shahvali, S., Rahiman, N., Jaafari, M.R. & Arabi, L. Targeting fibroblast activation protein (FAP): advances in CAR-T cell, antibody, and vaccine in cancer immunotherapy. Drug Deliv Transl Res 13, 2041–2056 (2023).

31. Gallant, J.P., et al. Identification and biophysical characterization of a novel domain-swapped camelid antibody specific for fentanyl. J Biol Chem 300, 107502 (2024).

32. Wei, W., Younis, M.H., Lan, X., Liu, J. & Cai, W. Single-Domain Antibody Theranostics on the Horizon. J Nucl Med 63, 1475–1479 (2022).

33. Klewinghaus, D., et al. Grabbing the Bull by Both Horns: Bovine Ultralong CDR-H3 Paratopes Enable Engineering of ’Almost Natural’ Common Light Chain Bispecific Antibodies Suitable For Effector Cell Redirection. Front Immunol 12, 801368 (2021).

34. Wang, Q., et al. Role of tumor microenvironment in cancer progression and therapeutic strategy. Cancer Med 12, 11149–11165 (2023).

35. de Visser, K.E. & Joyce, J.A. The evolving tumor microenvironment: From cancer initiation to metastatic outgrowth. Cancer Cell 41, 374–403 (2023).

36. Zou, B., et al. The Expression of FAP in Hepatocellular Carcinoma Cells is Induced by Hypoxia and Correlates with Poor Clinical Outcomes. J Cancer 9, 3278–3286 (2018).

37. Wikberg, M.L., et al. High intratumoral expression of fibroblast activation protein (FAP) in colon cancer is associated with poorer patient prognosis. Tumour Biol 34, 1013–1020 (2013).

38. Cohen, S.J., et al. Fibroblast activation protein and its relationship to clinical outcome in pancreatic adenocarcinoma. Pancreas 37, 154–158 (2008).

39. Hirmas, N., et al. Diagnostic Accuracy of (68)Ga-FAPI Versus (18)F-FDG PET in Patients with Various Malignancies. J Nucl Med 65, 372–378 (2024).

40. Liu, X., Liu, H., Gao, C. & Zeng, W. Comparison of (68)Ga-FAPI and (18)F-FDG PET/CT for the diagnosis of primary and metastatic lesions in abdominal and pelvic malignancies: A systematic review and meta-analysis. Front Oncol 13, 1093861 (2023).

41. Haberkorn, U., Altmann, A., Giesel, F.L. & Kratochwil, C. 1,090 Publications and 5 Years Later: Is FAP-Targeted Theranostics Really Happening? J Nucl Med 65, 1518–1520 (2024).

42. Hintz, H.M., Cowan, A.E., Shapovalova, M. & LeBeau, A.M. Development of a Cross-Reactive Monoclonal Antibody for Detecting the Tumor Stroma. Bioconjug Chem 30, 1466–1476 (2019).

43. Sun, X., et al. Beyond Small Molecules: Antibodies and Peptides for Fibroblast Activation Protein Targeting Radiopharmaceuticals. Pharmaceutics 16(2024).

44. Xu, J., et al. Screening and Preclinical Evaluation of Novel Radiolabeled Anti-Fibroblast Activation Protein-alpha Recombinant Antibodies. Cancer Biother Radiopharm 38, 726–737 (2023).

45. Dekempeneer, Y., et al. Preclinical Evaluation of a Radiotheranostic Single-Domain Antibody Against Fibroblast Activation Protein alpha. J Nucl Med 64, 1941–1948 (2023).

46. Wei, L., et al. Bamboo Shark as a Small Animal Model for Single Domain Antibody Production. Front Bioeng Biotechnol 9, 792111 (2021).

47. Hu, D., et al. Effective Optimization of Antibody Affinity by Phage Display Integrated with High-Throughput DNA Synthesis and Sequencing Technologies. PLoS One 10, e0129125 (2015).

48. Ribeiro, R., Moreira, J.N. & Goncalves, J. Development of a new affinity maturation protocol for the construction of an internalizing anti-nucleolin antibody library. Sci Rep 14, 10608 (2024).

49. Turner, K.B., et al. Next-Generation Sequencing of a Single Domain Antibody Repertoire Reveals Quality of Phage Display Selected Candidates. PLoS One 11, e0149393 (2016).

50. Miyazaki, N., et al. Isolation and characterization of antigen-specific alpaca (Lama pacos) VHH antibodies by biopanning followed by high-throughput sequencing. J Biochem 158, 205–215 (2015).

51. Yang, W., et al. Next-generation sequencing enables the discovery of more diverse positive clones from a phage-displayed antibody library. Exp Mol Med 49, e308 (2017).

52. Nannini, F., et al. Combining phage display with SMRTbell next-generation sequencing for the rapid discovery of functional scFv fragments. MAbs 13, 1864084 (2021).

53. Arras, P., et al. AI/ML combined with next-generation sequencing of VHH immune repertoires enables the rapid identification of de novo humanized and sequence-optimized single domain antibodies: a prospective case study. Front Mol Biosci 10, 1249247 (2023).

54. Erasmus, M.F., et al. Insights into next generation sequencing guided antibody selection strategies. Sci Rep 13, 18370 (2023).

55. Li, B., et al. Framework selection can influence pharmacokinetics of a humanized therapeutic antibody through differences in molecule charge. MAbs 6, 1255–1264 (2014).

56. Sanchez-Garrido, M.A., et al. Fibroblast activation protein (FAP) as a novel metabolic target. Mol Metab 5, 1015–1024 (2016).

57. Garin-Chesa, P., Old, L.J. & Rettig, W.J. Cell surface glycoprotein of reactive stromal fibroblasts as a potential antibody target in human epithelial cancers. Proc Natl Acad Sci U S A 87, 7235–7239 (1990).

58. Sum, E., et al. Fibroblast Activation Protein alpha-Targeted CD40 Agonism Abrogates Systemic Toxicity and Enables Administration of High Doses to Induce Effective Antitumor Immunity. Clin Cancer Res 27, 4036–4053 (2021).

59. Zielonka, S., et al. Structural insights and biomedical potential of IgNAR scaffolds from sharks. MAbs 7, 15–25 (2015).

60. Ubah, O.C., et al. Mechanisms of SARS-CoV-2 neutralization by shark variable new antigen receptors elucidated through X-ray crystallography. Nat Commun 12, 7325 (2021).

61. Limited, A. Positive AD-214 Phase I extension study results. Vol. 2024 Safety Announcement (Listcorp, www.listcorp.com, 2024).

62. Ubah, O.C., Porter, A.J. & Barelle, C.J. In Vitro ELISA and Cell-Based Assays Confirm the Low Immunogenicity of VNAR Therapeutic Constructs in a Mouse Model of Human RA: An Encouraging Milestone to Further Clinical Drug Development. J Immunol Res 2020, 7283239 (2020).

63. Steven, J., et al. In Vitro Maturation of a Humanized Shark VNAR Domain to Improve Its Biophysical Properties to Facilitate Clinical Development. Front Immunol 8, 1361 (2017).

64. Jung Min Kim, Robert M. Stroud & Craik, C.S. Rapid identification of recombinant Fabs that bind to membrane proteins. Methods 55(2011/12/01).

65. Brennen, W.N., et al. Mesenchymal stem cell infiltration during neoplastic transformation of the human prostate. Oncotarget 8(2017-04-21).

