## Supplemental Data for "Development of FAP-targeted theranostics discovered by next-generation sequencing-augmented mining of a novel immunized VNAR library"

### Supplemental Figure 1

| clone ID | clone frequency (%) | FW1 | CDR1 | FW2 | HV2 | FW3a | HV4 | FW3b | CDR3 | FW4 |
|---|---|---|---|---|---|---|---|---|---|---|
| H17 | 1.2 | ARVDQTPQTITKETGESLTINCVLR | DSNCALSS | TYWYRK | KSGSTNKESISKG | GRYVETV | NSGSK | SFSLRINDLTVEDSGTYRCKV | VYNWSEYDCGNSRFNYDV | YGDGTAVTVN |
| H12 | 1.2 | ARVDQTPQTITKETGESLTINCVLR | DRKCALSS | TYWYRK | KSGSTNEESIKKG | GRYVETV | NSGSK | SFSLRINDLTVEDSGTYRCNV | LMSWYGYPNEGLECWSDV | YGGGTVVTVN |
| H15 | 1.2 | ARVDQTPQTITKETGESLTINCVLR | DRKCALSS | TYWYRK | KSGSTNEESIKKG | GRYVETV | NSGSK | SFSLRINDLTVEDSGTYRCNV | LMSWYGYPNEGLECWSDV | YGDGTVVTVN |
| H13 | 1.2 | ARVDQTPQTITKETGESLTINCVL- | DSNCALSS | TYWYRK | KSGSTNEESISKG | GRYVETV | NSGSK | SFSLRINDLTVENSGTYRCNV | YVAGM----SPCLSWGDV | YGGGTVVTVN |
| H16 | 1.2 | ARVDQTPQTITKETGESLTINCVLR | DSNCALSS | TYWYRK | KSGSTNEESISKG | GRYVETV | NSGSK | SFSLRINDLTVEDSGMYRCNV | YVAGM----SPCLNWGDV | YGGGTVVTVN |
| H2 | 4.9 | ARVDQTPQTITKATGESLTINCVLR | DSNCALSS | TYWYRK | KSGSTNEESISKG | GRYVETV | ISGSK | SFSLRINDLTVEDSGTYRCNV | YVAGM----SPCLSWGDV | YGGGTVVTVN |
| H11 | 1.2 | ARVDQTPQTITKATGESLTINCVLR | DSNCALSS | TYWYRK | KSGSTNEESISKG | GRYVETV | ISGSK | SFSLRINDLTVEDSGTYRCNV | YVAGM----SPCLSWGDV | YGGGTAVTVN |
| H14 | 1.2 | ARVDQTPQTITKETGESLTINCVLR | DSNCALSS | TYWYRK | KSGSTNEESISKG | GRYVETV | NSGSK | SFSLRINDLTVEDSGTYRCNV | YVAGM----SPCLNWGDV | YGDGTAVTVN |
| H10 | 2.4 | ARVDQTPQTITKETGESLTINCVLR | DSNCALSS | TYWYRK | KSGSTNEESISKG | GRYVETV | NRGSK | SFSLRINDLTVEDSGTYRCNV | YVAGM----SPCLSWGDV | YGGGTVVTVN |
| H9 | 2.4 | ARVDQTPQTITKATGESLTINCVLR | DSNCALSS | TYWYRK | KSGSTNEESISKG | GRYVETV | ISGSK | SFSLRINDLTVEDSGTYRCNV | YVAGM----SPCLSWGDV | YGDGTAVTVN |
| H8 | 1.2 | ARVDQTPQTITKETGESLTINCVLR | DSNCALSS | TYWYRK | KSGSTNEESISKG | GRYVETV | ISGSK | SFSLRINDLTVEDSGTYRCNV | YVAGM----SPCLSWGDV | YGDGTAVTVN |
| H7 | 7.3 | ARVDQTPQTITKETGESLTINCVLR | DSNCALSS | TYWYRK | KSGSTNEESISKG | GRYVETV | NRGSK | SFSLRINDLTVEDSGTYRCNV | YVAGM----SPCLSWGDV | YGGGTAVTVN |
| H6 | 6.1 | ARVDQTPQTITKETGESLTINCVLR | DSNCALSS | TYWYRK | KSGSTNEESISKG | GRYVETV | NSGSK | SFSLRINDLTVEDSGTYRCNV | YVAGM----SPCLSWGDV | YGDGTVVTVN |
| H3 | 2.4 | ARVDQTPQTITKETGESLTINCVLR | DSNCALSS | TYWYRK | KSGSTNEESISKG | GRYVETV | NRGSK | SFSLRINDLTVEDSGTYRCNV | YVAGM----SPCLSWGDV | YGDGTAVTVN |
| H1 | 12.2 | ARVDQTPQTITKETGESLTINCVLR | DSNCALSS | TYWYRK | KSGSTNEESISKG | GRYVETV | NSGSK | SFSLRINDLTVEDSGTYRCNV | YVAGM----SPCLSWGDV | YGGGTAVTVN |
| H4 | 19.5 | ARVDQTPQTITKETGESLTINCVLR | DSNCALSS | TYWYRK | KSGSTNEESISKG | GRYVETV | NSGSK | SFSLRINDLTVEDSGTYRCNV | YVAGM----SPCLSWGDV | YGDGTAVTVN |
| H5 | 32.9 | ARVDQTPQTITKETGESLTINCVLR | DSNCALSS | TYWYRK | KSGSTNEESISKG | GRYVETV | NSGSK | SFSLRINDLTVEDSGTYRCNV | YVAGM----SPCLSWGDV | YGDGTVVTVN |

**Supplemental Figure 1, amino acid sequences of anti-FAP VNARs identified by biopanning.** Phagemids encoding anti-FAP VNARs were sequenced by Sanger sequencing, translated amino acid sequences are shown, along with their associated clone ID and the frequency of repeat sequences found among hit clones. Complementarity determining regions (CDR1, CDR3) and hypervariable loops (HV2, HV4) are depicted with black or gray shading, respectively.

**Supplemental Figure 2**

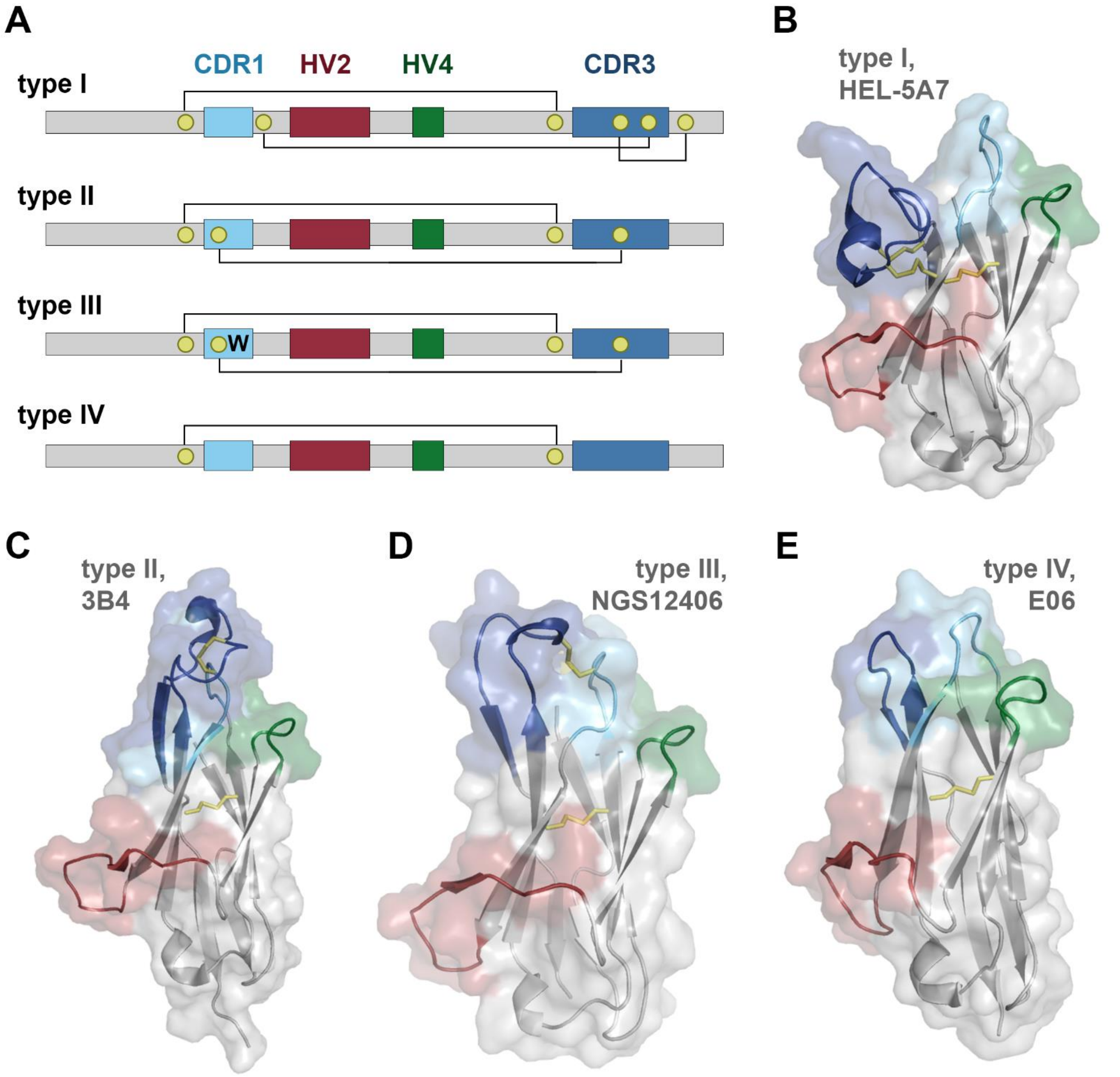


**Supplemental Figure 2, structural conformation of VNAR subtypes. A,** schematic of VNAR domain architecture. Framework regions (grey), complementarity determining region 1 (CDR1, cyan), hypervariable loop 2 (HV2, red), hypervariable loop 4 (HV4, green), complementarity determining region 3 (CDR3, blue), non-canonical cysteine residues (yellow), and conserved tryptophan residues (W) are shown. Disulfide bonds are illustrated with solid lines. **B-E,** cartoon and surface depictions of representative VNARs, colored as in (A), disulfide bonds are shown as yellow lines. B, type I VNAR, HEL-5A7 (PDB 1SQ2). C, type II VNAR, 3B4 (PDB 7SPO). D, Alphafold structure prediction of type III VNAR, NGS12406. E, type IV VNAR, E06 (PDB 4HGK).

**Supplemental Figure 3**

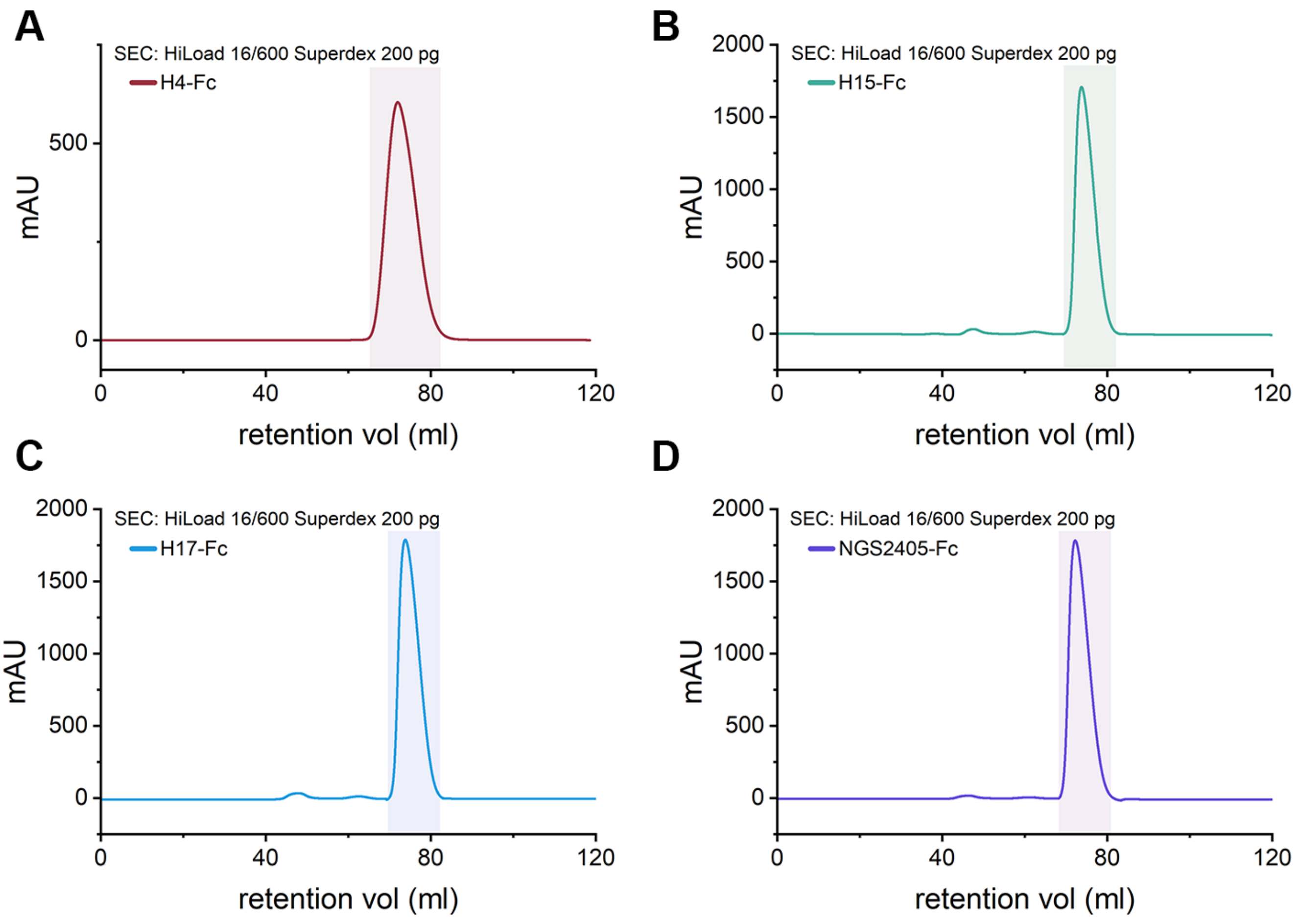


**Supplemental Figure 3, Size exclusion chromatography of VNAR-Fc constructs.** Anti-FAP VNAR-Fc constructs were purified by protein A affinity chromatography, size exclusion chromatography. Chromatograms of SEC of H4-Fc (A), H15-Fc (B), H17-Fc (C) and NGS2405-Fc (D) are shown./

**Supplemental Figure 4**

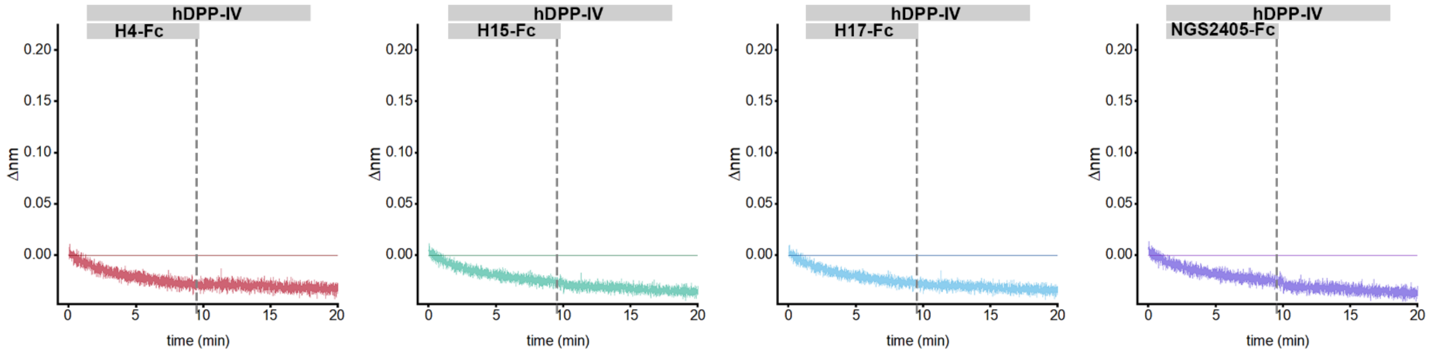


**Supplemental Figure 4, anti-FAP VNAR-Fc constructs do not bind to hDPP-IV.** Octet biosensors were loaded with biotinylated human DPP-IV and exposed to 2µM concentrations of either H4-Fc (red), H15-Fc (green), H17-Fc (blue), or NGS2405-Fc (purple).

**Supplemental Figure 5.**

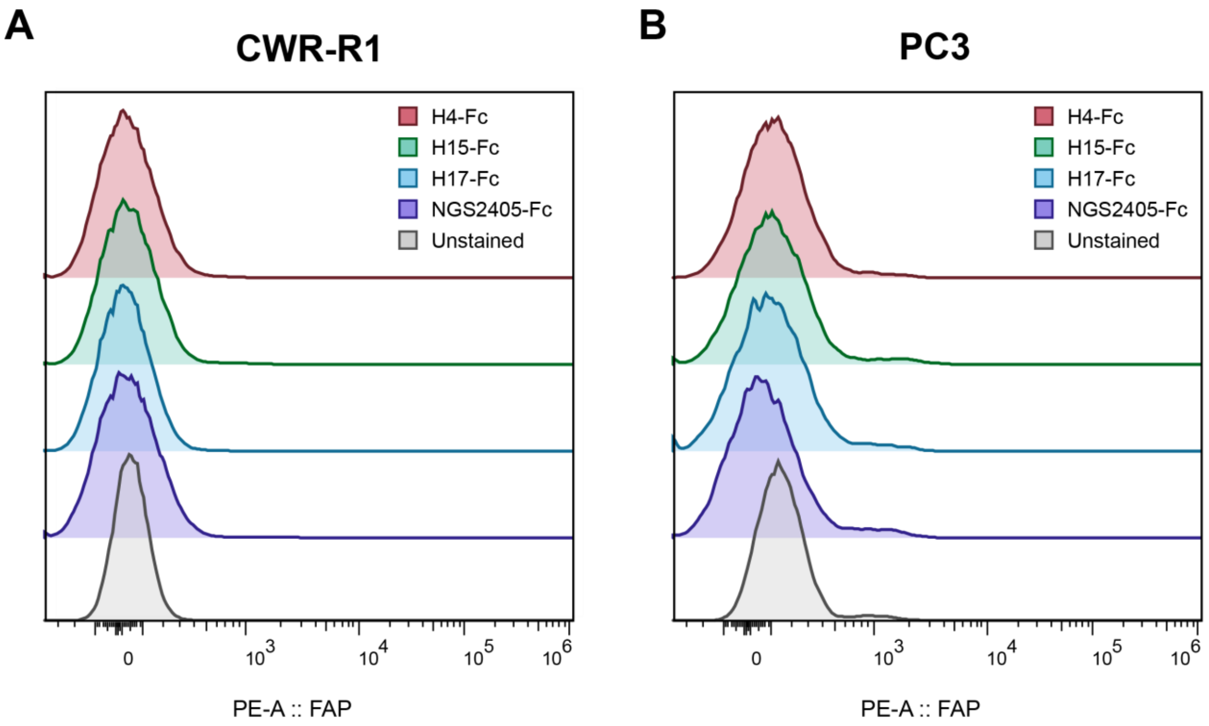


**Supplemental Figure 5, anti-FAP VNAR-Fc constructs fail to bind to FAP-negative CWR-R1 and PC-3 prostate cancer cells.** Assessing cellular binding of VNAR-Fc constructs to (A) CWR-R1 and (B) PC3 cell lines by flow cytometry. Cells were stained using a fixed concentration of VNAR-Fc antibodies (100 nM) and detected used a PE labeled anti-human Fc antibody. Samples were compared to an unstained cell control.

**Supplemental Figure 6**

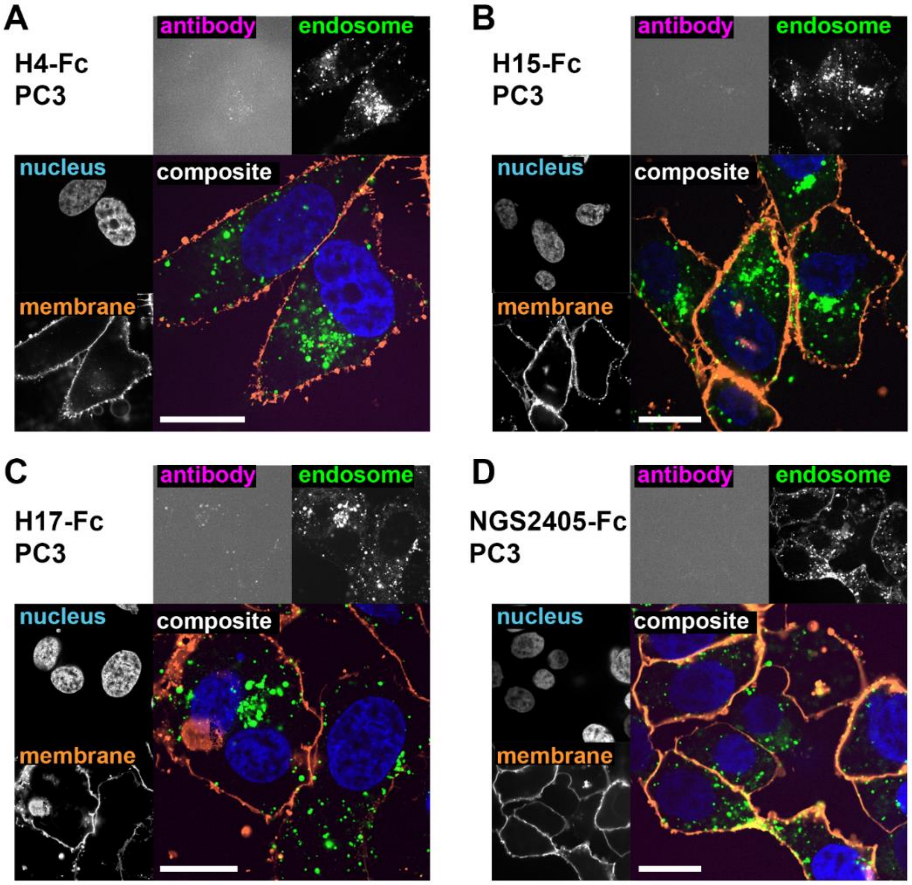


**Supplemental Figure 6, anti-FAP VNAR-Fc constructs fail to internalize into FAP-negative PC-3 prostate cancer cells.** Confocal microscopy images of PC-3 cells after incubation with H4-Fc-AF647 (A), H15-Fc-AF647 (C), H17-Fc-AF647 (E) or NGS2405-Fc-AF647 (G) for 1hr, using anti-FAP VNAR-Fc-AF647 (10nM) and fluorescein-dextran (50µg/ml). Single-channel images of fluorescein-labeled endosomes, Hoescht 33342-labeled nuclei, and CellBrite 555-labeled membranes are shown. Single-channel images of AF647 fluorescence are shown with high exposure to illustrate the lack of antibody internalization. Merged composite images depicting overlaid colorized fluorescent images are shown, scale bar represents 20µm.

**Supplemental Figure 7**

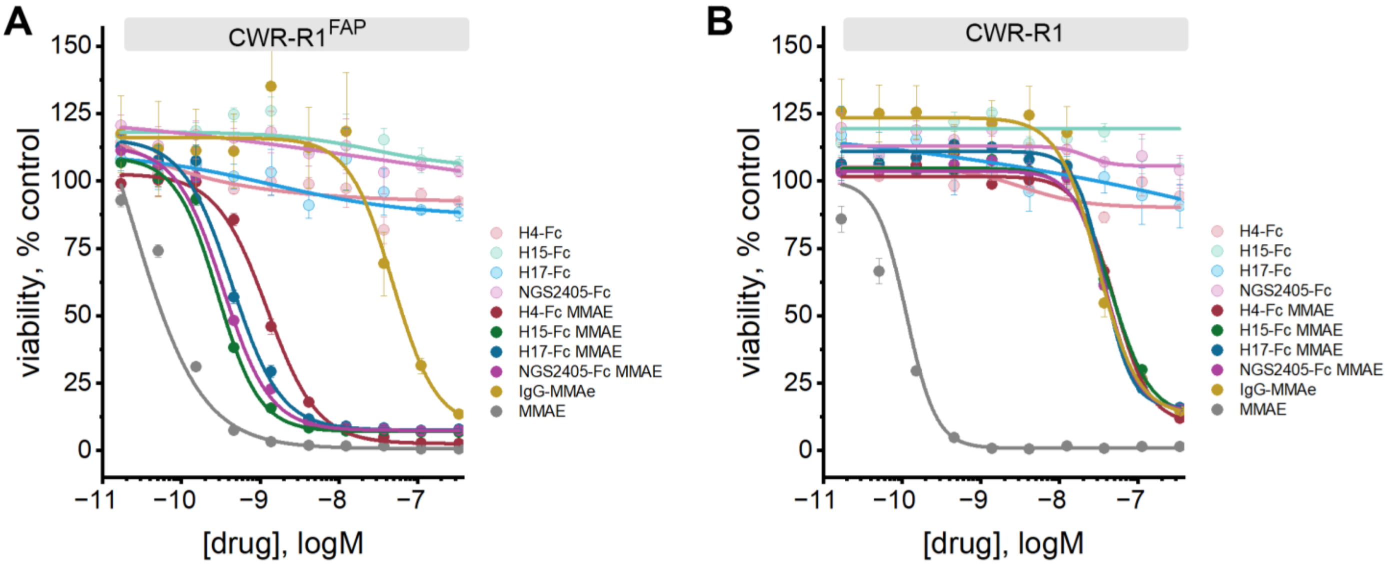


**Supplemental Figure 7, Anti-FAP VNAR-Fc-MMAE antibody-drug conjugates dose-dependently kill FAP-expressing cells.** Anti-FAP VNAR-Fcs site-specifically conjugated to a monomethyl auristatin E (MMAE) payload were tested for ability to kill CWR-R1$^{FAP}$ (A) and CWR-R1 (B) cells, as detected by measuring the NADPH reductive capacity in cells (CellTiter Blue) after incubation with serially diluted ADCs (300nM-0.03nM). Assays were conducted in parallel with parental unconjugated VNAR-Fc, a non-targeting isotype control VNAR-Fc-MMAE, and free MMAE drug. Data represents mean ± s.e.m. values from n=3 independent experiments.
